# Inhibition of IAPs induces programmed cell death and inflammatory signaling in patient-derived metastatic breast cancer organoids

**DOI:** 10.1101/2024.08.28.610103

**Authors:** Kaja Nicole Wächtershäuser, Jana V. Schneider, Alec Gessner, Geoffroy Andrieux, Ivan Kur, Nadine Duschek, Andreas Weigert, Melanie Boerries, Michael A. Rieger, Ernst H.K. Stelzer, Francesco Pampaloni, Sjoerd J.L. van Wijk

## Abstract

Breast cancer (BC) is the most common type of cancer among women worldwide and underlies relapse, disease progression and metastasis. Resistance to chemotherapy and programmed cell death (PCD), including apoptosis, strongly affects therapy success and remains a major challenge. Representative and translational models to understand, manipulate and cultivate advanced BC and to model PCD resistance are therefore urgently required. Smac mimetics are promising compounds to circumvent apoptosis resistance and are able to induce caspase-independent necroptosis, a lytic and inflammatory mode of PCD. Here, we apply primary, patient-derived human mammary organoids (hMOs) to investigate alternative forms of PCD to overcome apoptosis resistance. Using time lapse brightfield with immunofluorescent confocal microscopy, biochemistry and gene expression analysis, we demonstrate that Smac mimetics induce apoptosis in primary hMOs. By mimicking apoptosis resistance via caspase inhibition, hMOs undergo necroptosis, associated with expression and secretion of inflammatory mediators. Inhibition of linear ubiquitination by the LUBAC inhibitor HOIPIN-8 prevents necroptosis, as well as the expression and release of inflammatory mediators in hMOs. Our findings demonstrate that primary hMOs are effective models to model, study and manipulate PCD responses and inflammation in in primary BC organoids and open new therapeutic screening options for chemotherapy-resistant BC.

## Introduction

Breast cancer (BC) is the most commonly diagnosed type of cancer among women worldwide, leading to the most cancer-related deaths in females with steadily increasing incidence ^1,2^. The heterogeneity of BC, coupled with a rapid adaptation of tumor cells—involving metastasis and the development of resistance to chemotherapeutics and programmed cell death (PCD) mechanisms—significantly worsens the prognosis ^3,4^ and overall disease outcome ^5^.

To allow the investigation of multicellular tumor growth and PCD mechanisms, adult stem cell-derived organoids represent clinically and physiologically relevant experimental systems to study chemotherapy-treated and metastatic BC *in vitro* ^6,7^. These organoid cultures are established from patient tissues and maintain the characteristics of the original tumors ^8^. Organoids are grown in three-dimensional (3D) matrices that mimic the extracellular matrix (ECM) ^9,10^ and possess an inherent heterogeneity ^11^. The organization of organoids in biobanks makes patient-derived tumorigenic as well as healthy hMOs available for the research community ^8,12–15^. Of note, primary BC hMOs reflect the clinical response of patients prior to treatment and predict drug sensitivities ^6^. BC organoids allow personalized drug screening and target identification to overcome chemotherapy and PCD resistance in highly patient- and clinically relevant settings ^16^. In addition, the Food and Drug Administration (FDA) moved away from classical animal testing by approving the FDA Modernization Act 2.0 in 2022 and allows researchers to replace animal models with advanced *in vitro* models ^17^. Because of that, patient-derived organoids are bridging the gap between *in vitro* and *in vivo* studies ^18^.

Second mitochondria-derived activator of caspase (Smac) is the natural antagonist of inhibitor of apoptosis proteins (IAP) E3 ligases and induce protein clustering that leads to IAP auto-ubiquitination and proteasomal degradation ^19^. Cellular IAPs (cIAPs) as well as X-linked IAP (XIAP) are highly expressed in a wide variety of cancers and contribute to apoptosis resistance ^20^. Smac mimetics, including TL32711 (Birinapant), BV6, LCL-161 and ASTX660 can be used to induce PCD in various hematological and solid tumors ^21,22^ and multiple clinical trials have investigated the safety and efficacy of Smac mimetics as single agent or in combination with other compounds ^23^.

cIAPs, including cIAP1 and −2 as well as XIAP, typically mediate NF-ĸB, JNK and p38-dependent pro-survival responses via ubiquitination of cellular targets, including the receptor-interacting serine/threonine-protein kinase 1 (RIPK1), often downstream of death receptors, such as TNFR1 ^24^. Loss of RIPK1 ubiquitination upon IAP inhibition triggers the formation of pro-apoptotic complexes that activate caspase-8 to initiate caspase-3 and −7 cleavage and the execution of apoptosis ^25, 26^. Resistance to apoptosis occurs through mutations in extrinsic apoptosis pathways and overexpression of pro-survival proteins in other pro-apoptotic signaling pathways, including intrinsic apoptosis ^27,28^.

Caspase-independent forms of PCD, such as necroptosis, can overcome apoptosis resistance. Necroptosis is characterized by the auto- and trans-phosphorylation of RIPK1 and RIPK3, leading to the recruitment and phosphorylation of the mixed lineage kinase domain like pseudokinase (MLKL). Phosphorylated MLKL oligomerizes and translocates to the plasma membrane to cause membrane rupture and the release of cytokines, chemokines and damage-associated patterns (DAMPs) ^29^. Therefore, necroptosis can circumvent apoptosis resistance, but also triggers inflammatory responses in the cellular microenvironment to support anti-tumor immune responses.

Non-proteolytic linear ubiquitination, linked via methionine 1 (M1 poly-Ub), is an important checkpoint in survival and PCD responses and serves several cellular functions, including NF-ĸB activation. M1 poly-Ub is catalyzed by the linear ubiquitin chain assembly complex (LUBAC) ^30^ and M1 poly-Ub and LUBAC play important roles in cell survival, PCD and innate immunity ^31, 32^. LUBAC inhibition suppresses NF-ĸB signaling ^33^ and delays necroptotic cell death in several human cancer cell lines and healthy primary human pancreatic organoids ^34^.

To investigate PCD responses, apoptosis resistance and alternative forms of PCD in near-patient multicellular BC settings, we describe a translational experimental pipeline based on primary hMOs derived from metastatic lesions of BC patients. The hMOs exhibit donor-specific characteristics, including expansion rates, morphology and marker expression, thereby linking treatment outcome to BC heterogeneity. This setup allows for organoid- and single-cell-specific monitoring of morphology, apoptosis resistance and the induction of necroptosis as an alternative mode of BC cell killing. Furthermore, the role of LUBAC on cell death and PCD-associated inflammatory signaling was investigated. We provide a translational experimental platform to investigate PCD responses and resistance in primary hMOs and demonstrate that patient-derived organoids are a promising system for BC precision medicine.

## Methods

### hMO source and ethics

The human mammary organoids (hMOs) were obtained from HUB Organoid Technology (Utrecht, Netherlands) (donor #1 refers to organoid line HUB-C2-120, #2 to HUB-C2-152 and #3 to HUB-C2-123, as published in ^35^). The use of the primary hMOs has been approved by the medical ethical committee of the UMC Utrecht (METC UMCU; 12-427/C) under biobanking protocol HUB-Cancer TcBio#20-425. All patients signed informed consent forms and the experiments have been performed in accordance with the relevant guidelines and regulations.

### Chemicals and reagents

BV6 was kindly provided by Genentech Inc. (San Francisco, CA, USA), Birinapant, ASTX-660, LCL-161 and the pan-caspase inhibitor Emricasan were purchased from Selleckchem (Houston, TX, USA). Nec-1s and NSA from Merck KGaA (Darmstadt, Germany) and HOIPIN-8 from Axon Medchem LLC (Reston, VA, USA). All other chemicals were purchased from Carl Roth (Karlsruhe, Germany) or Merck KGaA (Darmstadt, Germany), unless stated otherwise.

### Maintenance of human mammary organoids (hMOs)

Primary hMOs were kept in an incubator at 37°C in a humidified atmosphere with 5% CO_2_ and were cultured, in adapted form, as described previously ^8,35^. hMOs from donor #1 and #2 were grown in modified Type I medium (Advanced Dulbecco’s Modified Eagles Medium with Nutrient Mixture F-12 (ThermoFisher Scientific, Waltham, MA, USA), 1 M HEPES (ThermoFisher Scientific), 1x GlutaMAX (ThermoFisher Scientific), 100 U/ml Penicillin/Streptomycin (ThermoFisher Scientific), 1.25 mM N-Acetylcyteine (Sigma, Schnelldorf, Germany), 500 nM A83-01 (R&D systems, Wiesbaden, Germany), 1x B27 without vitamin A (ThermoFisher Scientific), 5 µM Y-27632 (R&D systems), 10 mM nicotinamide (Sigma), 100 ng/ml recombinant Noggin-fc fusion protein (IpA Therapeutics, Fargo, ND, USA), R-spondin-3 conditioned medium at a concentration of 250 ng/ml R-spondin-3 (IpA Therapeutics), 5 ng/ml human EGF (PeproTech, London, UK), 50 µg/ml Primocin (InvivoGen, San Diego, CA, USA), 5 ng/ml FGF-7 (PeproTech), 37.5 ng/ml heregulin-β (PeproTech), 20 ng/ml FGF-10 (PeproTech), 500 nM SB202190 (Sigma). hMOs from donor #3 were maintained in modified Type II medium (Advanced Dulbecco’s Modified Eagles Medium with Nutrient Mixture F-12 (ThermoFisher Scientific, Waltham, MA, USA), 1 M HEPES (ThermoFisher Scientific), 1x GlutaMAX (ThermoFisher Scientific), 100 U/ml Penicillin/Streptomycin (ThermoFisher Scientific), 1.25 mM N-Acetylcyteine (Sigma, Schnelldorf, Germany), 500 nM A83-01 (R&D systems, Wiesbaden, Germany), 1x B27 without Vitamin A (ThermoFisher Scientific), 10 µM Y-27632 (R&D systems), 10 mM Nicotinamide (Sigma), 100 ng/ml recombinant Noggin-fc fusion protein (IpA Therapeutics, Fargo, ND, USA), 250 ng/ml recombinant R-spondin-3 (PeproTech), 500 ng/ml hydrocortisone (Sigma), 500 ng/ml human EGF (PeproTech), 50 µg/ml Primocin (InvivoGen), 10 µM Forskolin (R&D systems), 368 nM β-estradiol (Sigma), 0.25 nM WNT surrogate-Fc fusion protein (IpA Therapeutics), 5 ng/ml FGF-7 (PeproTech), 37.5 ng/ml heregulin-β (PeproTech), 40 ng/ml FGF-10 (PeproTech). Depending on proliferation rate and growth, organoids were passaged every seven to 21 days.

For passaging, hMOs were collected in 15 ml falcon tubes, washed with wash medium consisting of Dulbecco’s Modified Eagles Medium with GlutaMAX supplement (DMEM, ThermoFisher Scientific) supplemented with 0.1 % Albumin Fraction V (BSA, Carl Roth) and 100 U/ml Penicillin-Streptomycin (P/S, ThermoFisher Scientific) and spun down at 450 x g, 5 min and 8 °C. Washing was repeated using basal medium consisting of Advanced Dulbecco’s Modified Eagles Medium with Nutrient Mixture F-12 (ThermoFisher Scientific) supplemented with 1 M HEPES (ThermoFisher Scientific), 1x GlutaMAX (ThermoFisher Scientific) and 100 U/ml Penicillin-Streptomycin (P/S, ThermoFisher Scientific) and the remaining pellets were resuspended in 500 µl basal medium and 500 µl TrypLE™ Express Enzyme (ThermoFisher Scientific). hMOs were mechanically sheared into small fragments by pipetting, followed by washing in basal medium and subsequent resuspension in 20 % organoid medium with 80 % Cultrex Reduced Growth Factor BME, Type 2 (BME, R&D Systems) depending on the passaging ratio. BME droplets were placed in pre-warmed CELLSTAR® suspension culture plates (Greiner Bio-One GmbH, Frickenhausen, Germany) and solidified upside-down for 15 min in the incubator after which the droplets were overlaid with organoid medium.

### Time lapse imaging and live/dead assays

For time lapse imaging, hMOs were seeded in sterile CELLSTAR® 96-well suspension culture plates (Greiner Bio-One GmbH) in 5 µl BME droplets and overlaid with 100 µl organoid medium. Organoids were expanded for three to seven days and pre-treated with the indicated HOIPIN-8 concentrations for 30 min. Subsequently, the organoids were incubated with the Smac mimetics BV6, Birinapant, ASTX-660 or LCL-161, the RIPK1 inhibitor necrostatin-1s (Nec-1s), the MLKL inhibitor necrosulfonamide (NSA) or the pan-caspase inhibitor Emricasan (E) in the indicated concentrations. Plates were transferred to the Zeiss Z1 Axioimager widefield microscope (37 °C, 5 % CO_2_) and images were acquired every 30 min for a total period of 24 h with a 2 x 2 tiling and a Z-stack of 11 slices with 60 µm spacing.

For live/dead assays, hMOs were stained using 10 µg/ml propidium iodide (PI, Millipore Sigma), 0.5 µg/ml fluorescein diacetate (FDA, Millipore Sigma) and 200 µg/ml Hoechst33342 (ThermoFisher Scientific) for 30 min. The staining solution was removed and the droplets were washed with pre-warmed phosphate-buffered saline (PBS, ThermoFisher Scientific) prior to imaging in PBS using the same microscope settings as described above. Images were exported using the ZEN 2.6 lite software (Carl Zeiss Microscopy GmbH) and subsequently processed using the FiJi/ImageJ software.

### Viability measurements

hMOs were seeded and treated as described above. CellTiter-Glo® Luminescent Cell Viability (CTG, Promega) assays were performed after 24 h treatment by adding 10 µl CTG solution per well, followed by incubation for 30 min at RT in the dark. Luminescence was measured for 500 ms per well in an infinite M200 plate reader (Tecan, Männedorf, Switzerland). Technical triplicates were averaged per experiment and the background luminescence from control wells without CTG was subtracted from each value. The data were normalized against non-treated conditions and presented in bar plots with the standard deviation of three biological replicates using Origin 2023b Pro.

### hMO lysis and Western blotting

For lysis, hMOs were seeded in CELLSTAR® 24-well suspension culture plates (Greiner Bio-One GmbH) and expanded for up to ten days to reach high confluency. After treatment, two to six wells per condition were pooled and organoids were harvested in 15 ml falcon tubes, washed with wash medium, centrifuged at 600 x g, 5 min at 8 °C followed by washing with basal medium and PBS. Washes with PBS were repeated until no BME could be detected observed after which hMO pellets were stored at −20 °C until further processing. hMO pellets were thawed on ice and resuspended in RIPA lysis buffer (Pierce, ThermoFisher Scientific) supplemented with PhosSTOP™ (Hoffmann-LaRoche) and Protease Inhibitor Cocktail (Sigma) for 20 min on ice, followed by centrifugation at 15,000 x g and 4 °C for 15 min to remove cell debris. Protein concentrations were determined in cleared lysates using the BCA Protein Assay kit (Pierce, ThermoFisher Scientific) following the manufacturer’s protocol and using an infinite M200 plate reader (Tecan, Männedorf, Switzerland).

hMO lysates were diluted in water and denatured in 4x Laemmli buffer (ThermoFisher Scientific) with β-mercaptoethanol (Bio-Rad Laboratories, Hercules, CA, USA) to a final concentration of 1 mg/ml by incubating at 85 °C at 300 rpm for 10 min, followed by SDS-PAGE. The following antibodies were used in this study: α-Vinculin (V9131, Sigma), α-RIPK1 (610459, BD Biosciences), α-phospho-RIPK1 S166 (657465, Cell Signaling Technologies), α-RIPK3 (13526, Cell Signaling Technologies), α-phoshpo-RIPK3 S227 (ab209384, Abcam), α-MLKL (14993, Cell Signaling Technologies), α-phospho-MLKL S358 (91689, Cell Signaling Technologies), α-caspase-3 (9662S, Cell Signaling Technologies), α-cleaved-caspase-3 (9661, Cell Signaling Technologies), α-cIAP1 (AF8181, R&D Systems), α-cIAP2 (3130, Cell Signaling Technologies), α-XIAP (610716, BD Biosciences). The following horseradish peroxidase (HRP)-coupled secondary antibodies were used for detection using Pierce™ ECL Western Blotting-Substrate (ThermoFisher Scientific): goat α-rabbit-HRP (111-035-003, Jackson Immuno Research), goat α-mouse-HRP (115-035-003, Jackson Immuno Research), donkey α-goat-HRP (705-035-147, Jackson Immuno Research). Representative blots of at least two independent experiments are shown. If the samples of one experiment are detected on multiple Western blotting membranes, only one representative loading control is shown for clarity.

### RNA isolation, cDNA synthesis and quantitative real-time PCR

hMOs were seeded in CELLSTAR® 24-well suspension culture plates (Greiner Bio-One GmbH) with two wells per condition and grown to high confluency. After the indicated treatments, the culture medium was removed and hMO-containing BME droplets were resuspended in 500 µl TRIzol reagent (ThermoFisher Scientific) and collected at −20 °C.

For RNA extraction, 100 µl chloroform was added to each pellet, vigorously mixed and incubated for 3 min at RT, after which the phases were separated by centrifugation at 12,000 x g at 4 °C for 15 min. Approximately 250 µl of the aqueous phase were transferred into fresh tubes, mixed with 250 µl isopropanol and inverted five times. The tubes were incubated for 10 min at RT and centrifuged at 12,000 x g at 4 °C for 10 min. The supernatants were discarded and the RNA pellets were washed twice with 500 µl 75 % ethanol at 7,500 x g at 4 °C for 5 min. The pellets were subsequently dried and resuspended in 12 µl DNAse- and RNAse-free water (ThermoFisher Scientific). RNA concentrations were measured using NanoPhotometer® (Implen, Westlake Village, CA, USA) and 1.5 to 3 µg RNA were used for cDNA synthesis using the Maxima First Strand cDNA Synthesis Kit with dsDNAse (ThermoFisher Scientific) according to the manufacturer’s instructions.

For RT-qPCR, PowerTrack™ SYBR Green MasterMix (ThermoFisher Scientific) was used with a concentration of 5 ng of cDNA and 400 nM per primer per 10 µl reaction volume, using the CFX96 C1000 Touch qPCR system (BioRad). Data were normalized against *TBP, RPL13, 18S-rRNA* and *RPII* expression and relative gene expression levels were calculated using the ΔΔCt-method. The primers used in this study were designed using PrimerBlast ^36^ and are listed below. Primers were obtained from Eurofins (Hamburg, Germany) and biomers (Ulm, Germany). Primers for CASP1, CASP4 and CASP5 were acquired from RealTimePrimers (Elkins Park, PA, USA). CXCL1: forward AACCGAAGTCATAGCCACAC, reverse GTTGGATTTGTCACTGTTCAGC. CXCL10: forward CTGAGCCTACAGCAGAGGAAC, reverse GATGCAGGTACAGCGTACAGT. TNFα: forward ACAACCCTCAGACGCCACAT, reverse TCCTTTCCAGGGGAGAGAGG. ICAM1: forward CTTCCTCACCGTGTACTGGAC, reverse GGCAGCGTAGGGTAAGGTTC. 18S-rRNA: forward CGCAAATTACCCACTCCCG, reverse TTCCAATTACAGGGCCTCGAA, RPII: forward GCACCACGTCCAATGACAT, reverse GTGCGGCTGCTTCCATAA, TBP: forward TAAGAGAGCCACGAACCACG, reverse TTGTTGGTGGGTGAGCACAA, RPL13: forward AAGATCCGCAGACGTAAGGC, reverse GGACTCCGTGGACTTGTTCC,

### Immunofluorescence staining and confocal microscopy

hMOs were seeded in CELLSTAR® 24-well suspension culture plates (Greiner Bio-One GmbH) and grown to medium confluency. After treatment, hMOs were fixed using 4 % paraformaldehyde (PFA, Electron Microscopy Sciences) on ice for 30 min on a horizontal shaker and subsequently washed three times using ice-cold PBS. For immunostaining, organoids were transferred to 0.5 ml Eppendorf tubes permeabilized with 0.3 % Triton X-100 (Carl Roth) in PBS for 40 min at RT on a horizontal shaker. hMOs were washed three times 10 min with 100 mM glycine (Carl Roth) in PBS followed by three washing steps with PBS supplemented with 0.1 % Triton X-100 and 2 % P/S (PBS-T). Organoids are blocked in 0.2 % Triton X-100, 0.1 % Tween-20, 0.1 % BSA, 2 % P/S and 10 % donkey serum (Sigma) in PBS for at least 30 min at RT and 300 rpm. Primary antibody solutions were prepared in blocking solution and incubated with the organoids overnight at 37 °C and 300 rpm. The following primary antibodies were used in this study: α-E-cadherin (14472, Cell Signaling Technologies), α-Ki67 (ab16667, Abcam), α-Ki67 (9449, Cell Signaling Technologies), α-GATA-3 (ab199428, Abcam), α-CD49f (710209, ThermoFisher Scientific), α-cleaved caspase-3 (9661, Cell Signaling Technologies), α-phospho-RIPK1 (44590, Cell Signaling Technologies). The following day, organoids were washed once for 1 min followed by three times for 5 min with PBS-T and incubated with donkey α-mouse-AlexaFluor™568 (A10037, ThermoFisher Scientific) and donkey α-rabbit-AlexaFluor™647 (A32795, ThermoFisher Scientific) in blocking solution supplemented with 2 µg/ml Hoechst33342 (H1399, ThermoFisher Scientific) and 132 nM AlexaFluor™488 Phalloidin (A12379, ThermoFisher Scientific) for 2 h at 28 °C at 300 rpm. Prior to imaging, organoids were washed three times for 10 min in PBS-T and transferred onto a thin layer of CUBIC-2 (15 % w/w MilliQ H2O, 50 % w/w Sucrose (Carl Roth), 25 % w/w Urea (Carl Roth), 10 % w/v Triethanolamine (Carl Roth)) which was placed on a microscope slide (Epredia, Portsmouth, NH, USA). A cover glass was placed on top and the organoids were imaged using a Zeiss LSM780 confocal microscope with a Plan-Apochromat 20x/0.8 M27 objective, unless indicated otherwise. Image processing was performed using Zeiss ZEN 3.8 and FiJi/ImageJ software.

### Cytometric Bead Array (CBA) assays

Cytokine and chemokine secretion of hMOs was determined by seeding organoids in 10 µl drops in 96-well plates in 100 µl medium to high confluency. Organoids were treated in 70 µl medium for 24 h, after which supernatants were collected and stored at −80°C. Assays were performed with the BD® Cytometric Bead Array (CBA) assay (Becton Dickinson (BD), Heidelberg, Germany), according to the manufacturer’s instructions. Briefly, a master-mix of cytokine capture beads and CBA buffer was prepared and 25 µl were added to FACS tubes and mixed with 25 µl of the hMO supernatants, followed by vortexing and incubation for 1 h at RT. 25 µl of the detection reagent master-mix was added to each FACS tube and incubated for another 2 h at RT in the dark. Subsequently, samples were washed with 1 ml FACS Flow™ sheath fluid (BD) and beads were pelleted by centrifugation at 200 x g for 5 min at RT. Supernatants were discarded and the pellets were resuspended in 150 µl FACS Flow™ sheath fluid, after which cytokine- and chemokine levels were determined using FACSymphony™ A5 SE (BD). Analysis was performed with BD FACS DIVA Software (BD).

### Statistical analysis

Statistical significance was determined using an unpaired, two-tailed student’s t-test assuming unequal variances. p values < 0.05 are considered significant and depicted as follows: p ≤ 0.05: *, p ≤ 0.01: **, p ≤ 0.005: ***, ns: not significant.

## Results

### Establishing an experimental pipeline to monitor primary hMOs

To bridge the gaps between *in vitro* and *in vivo* models of breast cancer and to reduce the use of animal models, we investigated if primary, patient-derived hMOs can be applied as preclinical models to study multicellular tumor growth and sensitivities towards PCD. This is facilitated by using advanced, chemotherapy-treated and metastatic human breast cancer mammary organoids (hMOs), that maintain the original tumor characteristics, and which are derived from adult stem cells and obtained from patient tissues. Here, using these primary hMOs, we aimed to setup an experimental system to systematically study PCD sensitivity and inflammation. Three hMOs lines were obtained from biobanking and stable cultures were established in a chemically defined medium that mimics the tumor microenvironment. hMO growth and expansion were monitored optically using brightfield imaging as well as immunofluorescence (IF) and confocal microscopy.

Consistent with the clinical situation ^8^, the three hMO lines exhibit distinct morphological features and show different expansion rates during routine culturing (data not shown). hMOs derived from donor #1 and donor #3 presented a compact, round phenotype, whereas the BC organoids of donor #2 proliferated in a “grape-like” morphology. All hMOs contained Ki67-positive nuclei, but more Ki67-positive cells could be detected in the less dense growing hMOs from donor #2 (Figure 1A), indicating that Ki67 levels reflected expansion rates. hMOs from donor #1 and #3 expressed E-cadherin, whereas no E-cadherin could be detected in hMOs from donor #2 (Figure 1B), suggesting a potential link between E-cadherin expression and proliferation.

**Figure 1:**
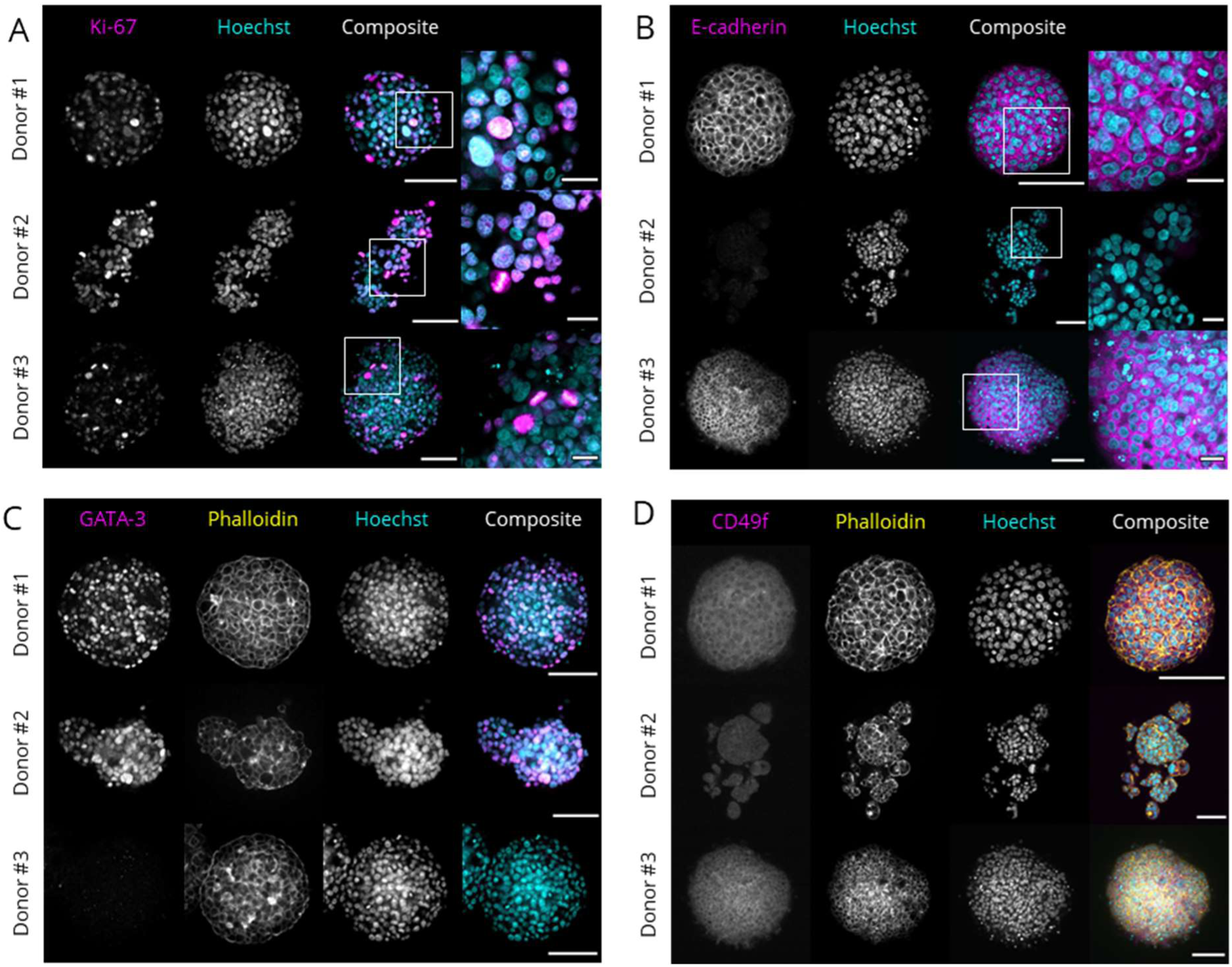
Establishing an experimental pipeline to monitor primary hMOs. hMOs from three independent donors were grown until confluency in growth medium, fixed and subjected to immunofluorescent staining against Ki-67 (A), E-cadherin (B), GATA-3 (C) and CD49f (D) (magenta) and counterstained using Hoechst33342 (cyan) and AF488-phalloidin (yellow). Organoids were cleared with CUBIC-2 and imaged using a Zeiss LSM780 confocal microscope. Scale bar: 100 µm, enlarged image: 25 µm.

To further investigate donor-specific differences in mammary marker expression, GATA-3 and CD49f expression were evaluated using IF confocal microscopy (Figure 1C & D). GATA-3 is a marker for luminal cells ^37^ and highly expressed in luminal A BC ^38^. The highest levels of GATA-3 expression could be detected in hMOs from donor #2, and to a lower extent in donor #1 (Figure 1C) and showed nuclear localization. Donor #3 exhibited almost no GATA3-positive cells. CD49f, also referred to as integrin- α6, is a biomarker for breast cancer ^39^ and is expressed in hMOs from all three donors (Figure 1D).

In summary, luminal A and B hMOs from three independent patients were stably cultured and expanded over extended periods and demonstrated proliferative phenotypes, with highly donor-specific differences in expansion rate and tumor marker expression.

### Smac mimetics induce cIAP1/2 degradation and donor-specific loss of cell viability in primary hMOs

To monitor Smac mimetic-induced changes in hMO viability, stably growing hMOs were incubated with the Smac mimetics BV6, Birinapant, ASTX-660 and LCL-161 for 24 h, followed by the analysis of cell viability. In all three donors, BV6 induced the strongest reduction in cell viability, whereas Birinapant, ASTX-660 and LCL-161 affect hMO viability in a highly donor-specific manner (Figure 2A). Western Blot analysis of cIAP1, cIAP2 and XIAP expression in non-treated, DMSO or Smac mimetic-treated hMOs revealed compound-induced degradation of cIAP1 in lysates of all hMOs (Figure 2B). Clear cIAP2 expression was only detected in hMOs of donor #3 and was strongly reduced with all Smac mimetics tested. The hMOs from all donors expressed XIAP and potent loss of XIAP was observed by BV6 and to a lesser extent with the other Smac mimetics (Figure 2B). All donors expressed caspase-3 and Smac mimetics induced caspase-3 cleavage in a donor-specific manner, suggesting the induction of apoptosis (Figure 2B).

**Figure 2:**
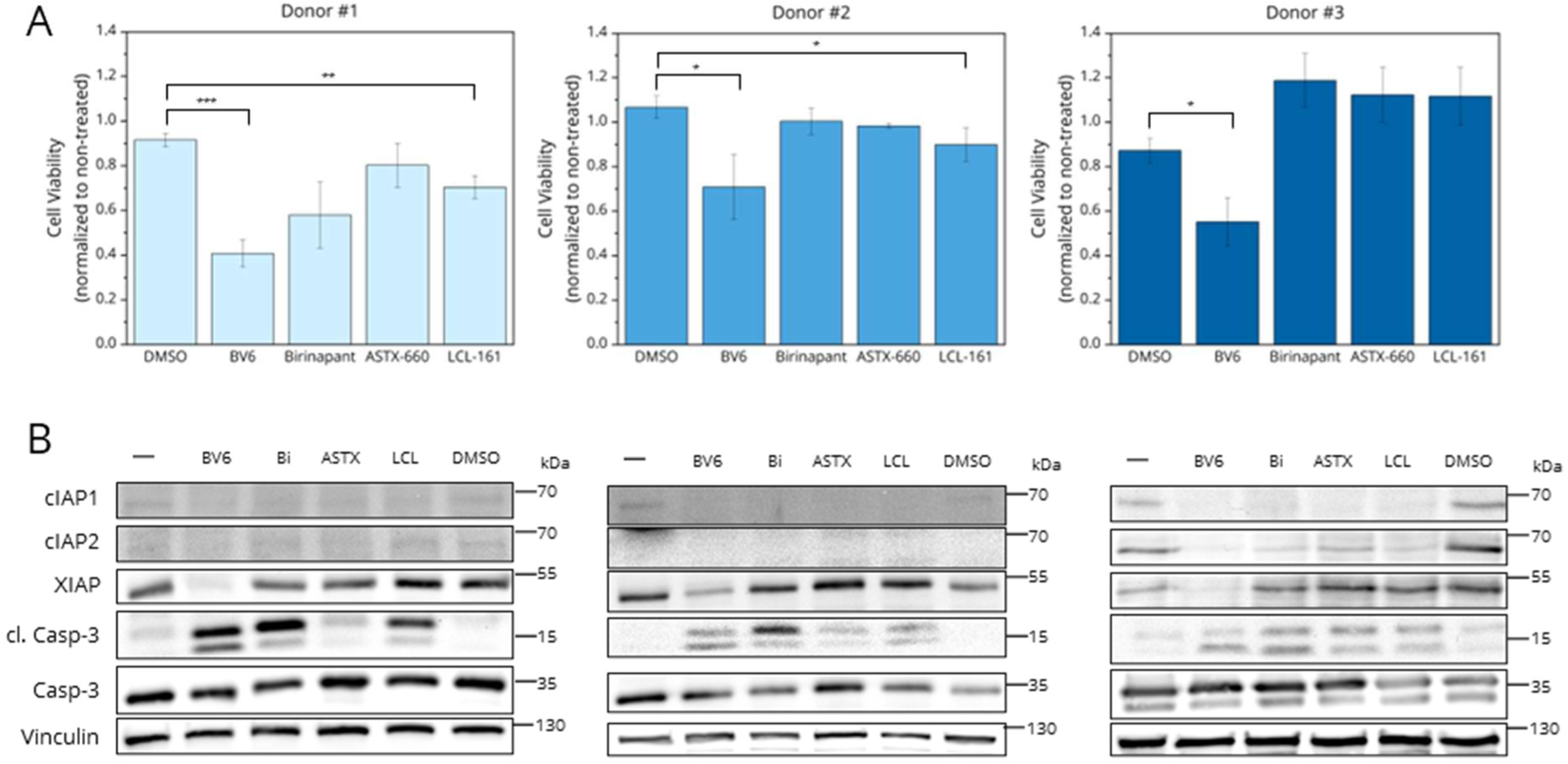
Smac mimetics induce cIAP1/2 degradation and donor-specific loss of cell viability in primary hMOs. (A) hMOs were grown for three to seven days in growth medium followed by 24 h treatment with 10 µM of BV6, Birinapant (Bi), ASTX-660 (ASTX) and LCL-161 (LCL), respectively, or DMSO as vehicle control. Subsequently, CellTiter-Glo® viability assays were performed and the values were normalized to non-treated controls. Error bars represent standard deviation. * p < 0.05, ** p< 0.01, *** p < 0.005. (B) hMOs, treated as in (A) were harvested and Western blotting was performed with antibodies recognizing cIAP1, cIAP2, XIAP as well as total and cleaved caspase-3. Vinculin served as loading control. Representative blots of three independent experiments are shown.

### Smac-mimetics induce programmed cell death in primary hMOs

Since BV6 and Birinapant demonstrated the strongest effects on hMO cell viability and since Birinapant is already being tested in clinical trials, we further focused on Birinapant and BV6-induced PCD in primary hMOs. hMOs were treated with the compounds, or DMSO as control, and subjected to time lapse brightfield imaging. Treatment of primary hMOs with Birinapant induced morphological alterations (Figure 3A, Supplementary Figure 1A & C). Morphologically, Smac mimetic-treated hMOs adapt a looser morphology and especially the hMOs from donor #1 and donor #3 lost their compact phenotypes (Figure 3A & B, Supplementary Figure 1C & D). After 12 h, individual cells started to emerge from the compact organoids and the formation of apoptotic bodies could be observed. Subsequent live-dead assays to distinguish between viable cells using fluorescein-diacetate (FDA, green), dead cells using propidium iodide (PI, red) and total cells with Hoechst33342 (Hoechst, blue) revealed increased cell death Birinapant-treated hMOs compared to non-treated organoids (Figure 3B, Supplementary Figure 1B & D). Furthermore, Birinapant-treated hMOs stained positive for cleaved caspase-3, indicating prominent induction of apoptosis (Figure 3C). Individual cells within the hMOs appeared as more rounded as shown upon phalloidin-based actin staining (Figure 3C, Supplementary Figure 2 A & B).

**Figure 3:**
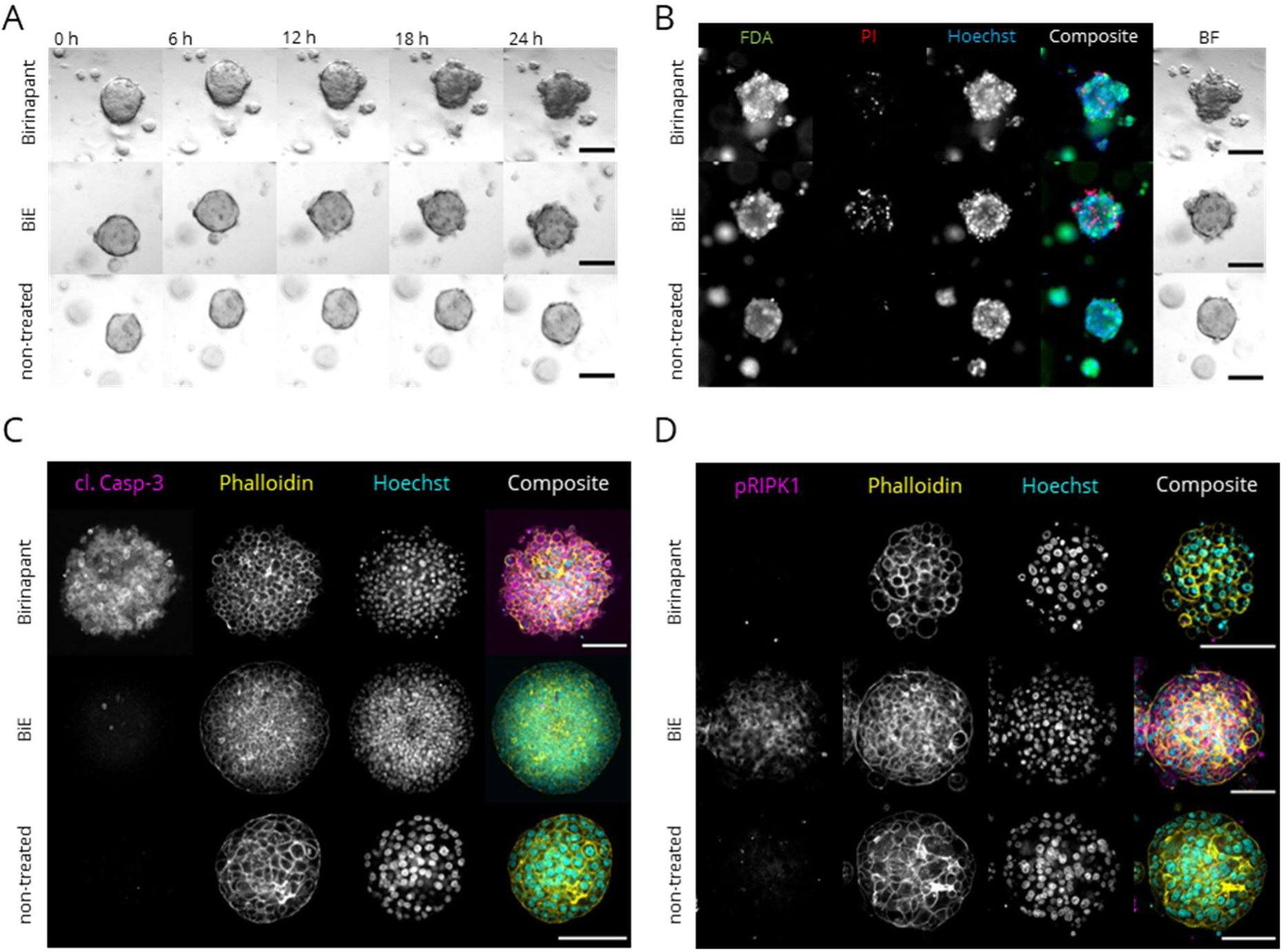
Smac-mimetics induce programmed cell death in primary hMOs. (A) hMOs were grown for three to seven days followed by 24 h treatment with 10 µM Birinapant and in combination with 10 µM Emricasan (BiE). Brightfield (BF) time-lapse imaging was performed using the Zeiss Z1 Axioimager widefield microscope with 30 min intervals. (B) hMOs from (A) after live imaging were stained using fluorescein-diacetate (FDA, viable cells, green), propidium iodide (PI, dead cells, red) and Hoechst33342 (Hoechst, all nuclei, blue). (C & D) hMOs, treated as in (A), were fixed and subjected to immunofluorescence staining against cleaved caspase-3 (cl. Casp-3, magenta) and S166-phosphorylated RIPK1 (pRIPK1, magenta) and counterstained using AF488-phalloidin (yellow) and Hoechst33342 (cyan). Organoids were cleared with CUBIC-2 and imaged using a Zeiss LSM780 confocal microscope. (A-D) Representative images of donor #1, scale bar: 100 µm.

BV6 treatment induced similar effects to Birinapant, but the morphological changes appeared already after 6 h (Supplementary Figure 3 A, C & E) and resulted in a stronger PI signal indicating higher numbers of dead cells (Supplementary Figure 3B, D & F). IF staining against cleaved caspase-3 revealed areas of higher apoptosis activation within the organoid (Supplementary Figure 4 A-C). Especially, the cells that migrated out of the compact organoid in hMOs from donor #1 showed high levels of caspase-3 cleavage (Supplementary Figure 4A).

To model apoptosis resistance, which is commonly observed in relapsed or metastasized breast cancer, and to investigate if primary hMOs are susceptible to necroptosis induction, caspases were inhibited by the pan-caspase inhibitor Emricasan. Birinapant- and Emricasan-cotreated hMOs appeared as induced spheres, indicating cell swelling after 18 h (Figure 3A, Supplementary Figure 1 A & C) which is associated with necroptosis ^40^. Birinapant- and Emricasan-cotreated hMOs indicated more prominent cell death, indicated by more PI-positive cells (Figure 3B, Supplementary Figure 1 B & D). In Birinapant- and Emricasan-cotreated hMOs, no cleaved caspase-3 could be observed, pointing towards efficient blockage of apoptosis (Figure 3C). In contrast, prominent RIPK1 phosphorylation associated with necroptosis was detected and exhibited a homogenous distribution throughout the necroptotic organoids (Figure 3D). Of note, RIPK1 phosphorylation was absent in untreated hMOs and hMOs treated with Birinapant alone (Figure 3D, Supplementary Figure 2). Necroptotic hMOs showed a more compact morphology similar to non-treated hMOs with individual cells undergoing swelling and expulsion from the organoid (Figure 3D, Supplementary Figure 2).

hMO cotreatment with BV6 and Emricasan also induced potent cell swelling over 24 h and resulted in increases in PI-positive cells. In line with this, FDA signals pointing towards viable cells were greatly reduced compared to DMSO control-treated organoids (Supplementary Figure 3). Induction of necroptosis could be confirmed by IF staining of phosphorylated RIPK1 upon BV6-Emricasan co-treatment, but not in hMOs incubated with BV6 only (Supplementary Figure 4).

Taken together, these data suggest that cell death can be differentially induced and monitored in primary hMOs from multiple independent donors and that caspase inhibition can be applied to mimic the induction of apoptosis and necroptosis.

### Necroptosis and LUBAC inhibitors rescue Birinapant- and Emricasan-induced PCD in primary hMOs

To confirm the onset and occurrence of necroptosis, we applied the RIPK1 inhibitor necrostatin-1s (Nec-1s), the MLKL inhibitor necrosulfonamide (NSA) and the LUBAC inhibitor HOIPIN-8 (H) on Birinapant- and Emricasan-treated hMOs. Using these inhibitors, no morphological changes associated with PCD could be observed in the hMOs during time lapse imaging (Figure 4A, Supplementary Figure 5 A & C) and no cell death could be observed in the fluorescent live/dead assay (Figure 4 B, Supplementary Figure 5 B & D). Moreover, inhibition of RIPK1, MLKL and to a lesser extent also LUBAC, rescued hMO cell viability (Figure 4 C, Supplementary Figure 5 E), although hMOs of donor #3 showed no reduction in cell viability when treated with Birinapant and Emricasan, which did not further change upon incubation with inhibitors (Supplementary Figure 5 G). Western Blot analysis revealed prominent induction of RIPK1, RIPK3 and MLKL phosphorylation in BiE-treated hMOs, which was absent in untreated, DMSO or Bi treated hMOs (Figure 4 D, Supplementary Figure 5 F & H). Inhibition of RIPK1 inhibited phosphorylation of RIPK1 itself, RIPK3 and MLKL, whereas NSA did not affect RIPK1, RIPK3 and MLKL phosphorylation but prevents necroptosis by interfering with MLKL membrane translocation ^41,42^. Interestingly, inhibition of LUBAC did not affect phosphorylation patterns, underscoring the initial observations that HOIPIN inhibits necroptosis downstream of MLKL phosphorylation^34^. These data suggest that necroptosis in primary hMOs depends on activation of RIPK1, RIPK3 and MLKL and that inhibition of LUBAC prevents necroptosis in primary hMOs. Similar findings are detected with BV6 (Supplementary Figures 6 and 7). Of note, BV6- and Emricasan-induced cell death could be inhibited with Nec-1s, NSA and HOIPIN-8, but cell viability was only rescued in hMOs derived from donor #1 and #2 (Supplementary Figure 7 E).

**Figure 4:**
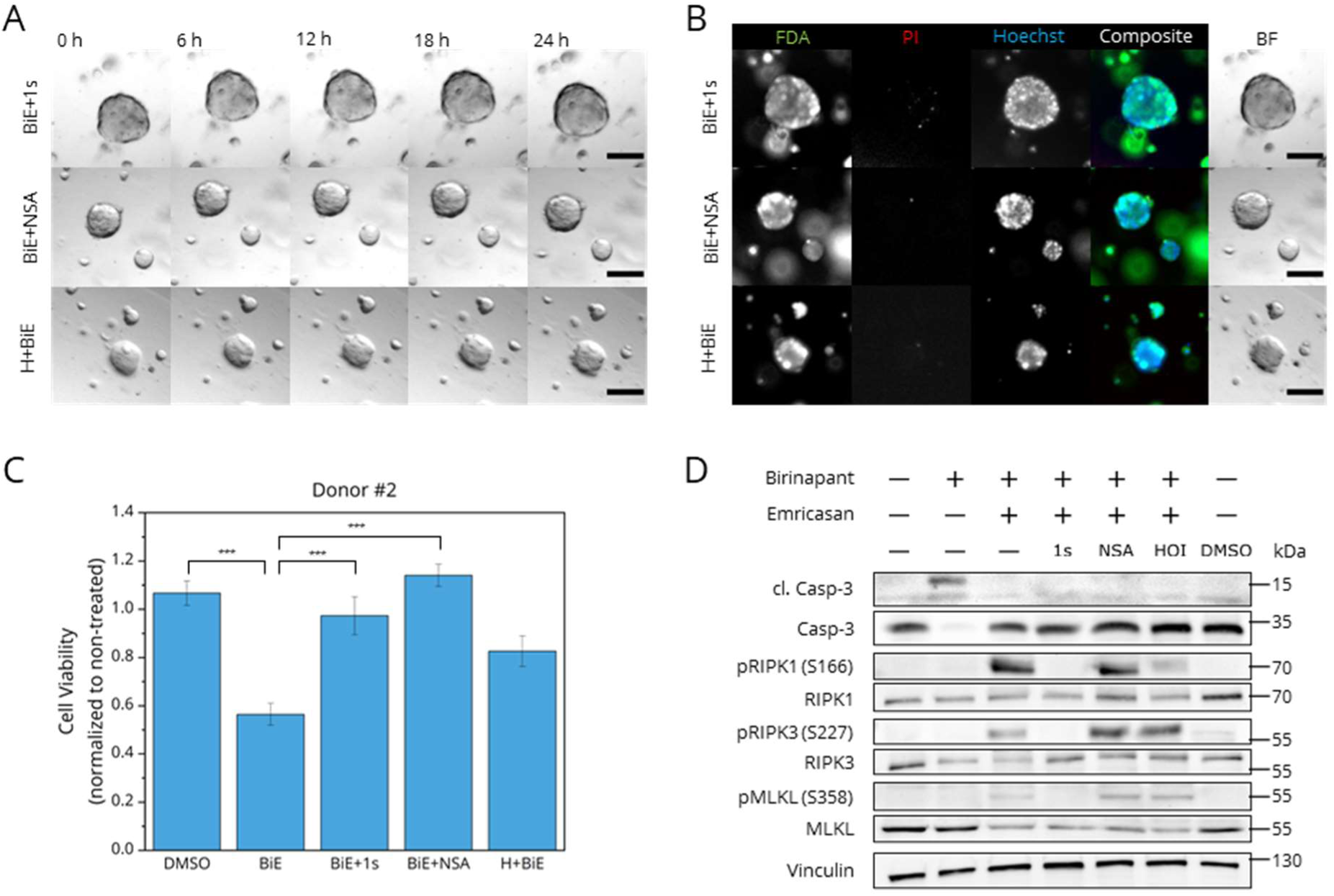
Necroptosis and LUBAC inhibitors rescue Birinapant- and Emricasan-induced PCD in primary hMOs. (A) hMOs were grown for three to seven days and pre-treated with 30 µM HOIPIN-8 (H, HOI) for 30 min followed by treatment with 10 µM Birinapant and in combination with 10 µM Emricasan (BiE), 30 µM necrostatin-1s (1s) and 10 µM necrosulfonamide (NSA) as indicated. Subsequently, hMOs were subjected to brightfield (BF) time lapse imaging using the Zeiss Z1 Axioimager widefield microscope with 30 min intervals. (B) hMOs from (A) after live imaging were stained using fluorescein-diacetate (FDA, viable cells, green), propidium iodide (PI, dead cells, red) and Hoechst33342 (Hoechst, all nuclei, blue). Representative images of donor #1, scale bars: 100 µm. (C) CellTiter-Glo® viability assays were performed on hMOs from donor #2 treated as in A. Values were normalized to non-treated controls. Error bars represent standard deviation. *** p < 0.005. (D) hMOs, treated as in A were harvested and Western blotting was performed with antibodies recognizing phosphorylated and total RIPK1, RIPK3 and MLKL. Vinculin served as loading control. Representative blots of three independent experiments are shown.

### The induction of necroptosis triggers the release of inflammatory mediators in primary hMOs

Necroptosis is highly inflammatory, not only via the release of DAMPs upon membrane rupture, but also by secretion of inflammatory mediators, including cytokines and chemokines. Consistent with the induction of necroptosis, increased TNF and CXCL10 mRNA expression (Figure 5 A & C) was found in Birinapant- and Emricasan-treated hMOs from all three donors. Overall, an upregulation of TNF and CXCL10 mRNA in Birinapant- and Emricasan-treated hMOs could be observed, compared to Birinapant-treated hMOs. Cotreatment with HOIPIN-8 decreases TNF and CXCL10 mRNA expression (Figure 5 A & C). Next, we investigated the secretion of TNF-α and IP-10 using CBA FACS-based analysis. Intriguingly, only BiE-induced hMOs from donor #2 upregulated TNF-α secretion, whereas TNF-α was induced by Bi in donor #1 and hMOs from donor #3 were non-responsive to either Birinapant or BiE (Figure 5 B). Cotreatment with HOIPIN-8 reduced BiE-induced TNF-α and IP-10 secretion, consistent with mRNA expression and previous findings ^34^ (Figure 5 A-D). Similar responses were observed for secretion of pro-inflammatory IL-6 and IL-8 (Supplementary Figure 8A).

**Figure 5:**
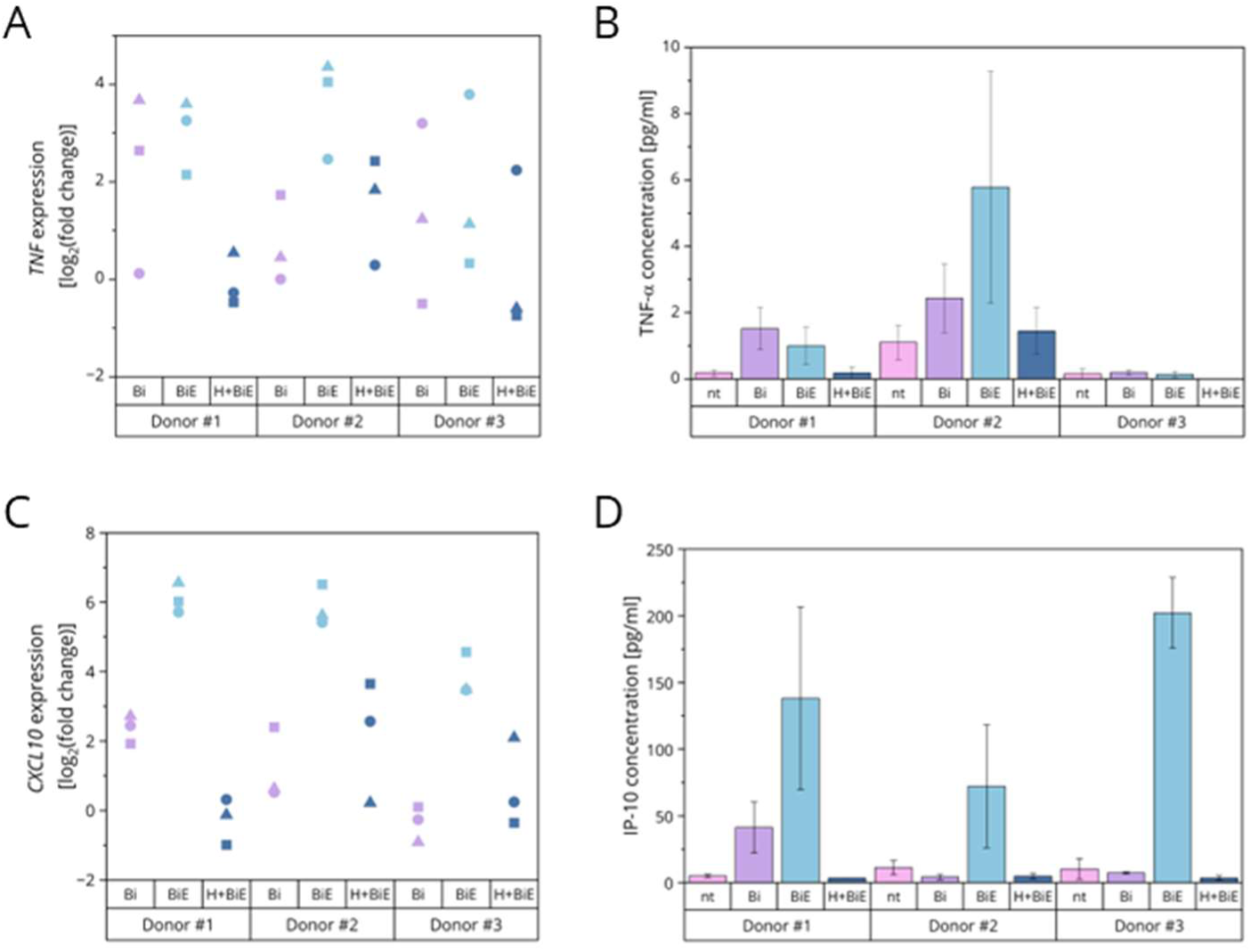
The induction of necroptosis triggers the expression and secretion of inflammatory mediators in primary hMOs. (A) mRNA expression levels of *TNF* in the indicated hMOs grown for three to seven days, pre-treated with 30 µM HOIPIN-8 for 30 min followed by treatment with 10 µM Birinapant (Bi) and in combination with 10 µM Emricasan (BiE). Gene expression was normalized against non-treated conditions and *RPII*, *18S*, *TBP* and *RPL13* mRNA expression and is presented as log_2_(fold change). (B) The indicated hMOs were grown and treated as in (A) and supernatants were subjected to FACS-based CBA assays to quantify secretion of TNF-α. Mean and standard deviation of three independent experiments are shown. (C) Same as (A), but *CXCL10* mRNA levels were quantified. (D) Same as (B), but IP-10 secretion was quantified.

Inflammatory caspases-1, −4 and −5 are linked to pyroptosis, aninflammatory form of PCD ^43,44^. Particularly, caspase-1 is known to cleave pro-IL-1β and to modulate inflammation ^45,46^. Intriguingly, in donor #1 and #2, *CASP1* expression was higher in BiE-treated organoids compared to Bi-treated organoids, which showed similar *CASP1* expression levels to non-treated control. HOIPIN-8 cotreatment led to a slight reduction in *CASP1* expression, following the same trend observed for *TNF* and *CXCL10* expression. The expression levels of the other caspases increased upon induction of necroptosis in donors #1 and #2, but cotreatment with HOIPIN-8 did not lead to the reduction as in *CASP1* expression. Donor #3 had similar caspase expression levels in all conditions (Supplementary Figure 8B). Even though caspases were inhibited, when Emricasan was applied, the transcript expression levels were increased. Contrary to caspases-4 and −5 ^45^, caspase-1 is part of the PANoptosome and drives apoptosis, necroptosis and pyroptosis ^47–49^. In agreement with our previous findings that HOIPIN-8 attenuates necroptosis, *CASP1* expression was downregulated by cotreatment with HOIPIN-8.

Similar effects could be observed with BV6 (Supplementary Figure 9).

In summary, our findings confirm the establishment of an experimental pipeline to study programmed cell death responses, sensitivities and resistance in primary, patient-derived organoids. Differential sensitivities against apoptosis and necroptosis could be modelled and necroptosis-associated inflammatory signaling was determined Furthermore, LUBAC inhibition by HOIPIN-8 reversed inflammatory and cell death responses linked to necroptosis. The combination of primary tumorigenic organoids, cell death and inflammation opens novel possibilities to implement organoids in translational personalized and precision medicine approaches.

## Discussion

Breast cancer (BC) is the most common form of invasive cancer in women with increasing incidence. Despite advances in treatment options, relapse rates remain high due to primary and acquired chemoresistance [3]. Loss-of-function mutations and upregulation of factors that counteract apoptotic PCD pathways critically underlie chemoresistance and promote metastasis [3]. In addition, adequate PCD programs affect the microenvironment, local and organismal responses and tissue damage upon tumor formation [4]. The activation of alternative modes of PCD, such as necroptosis, might therefore be highly attractive to overcome acquired resistance against chemotherapy and apoptosis in BC. Unfortunately, the study of necroptosis in human cancer settings is restricted only to a handful of necroptosis-proficient human tumor cell lines, that are still limited due to cell type-specific RIPK1, RIPK3 and MLKL expression. Up till now, primary and multi-cellular models that recapitulate complex cell-cell contacts, 3D interactions and the physiological tissue-like cellular microenvironment to investigate the molecular mechanisms of necroptosis in human materials are not available. Here, we describe a translational pipeline to model, mimic and manipulate PCD sensitivities and associated inflammatory signaling in primary, patient-derived hMOs.

We established long-term, stable cultures of primary hMOs from independent luminal A (donor #2 and #3) and B (donor #1) BC patients. These hMOs differ morphologically from compact and round, to grape-like and relatively fragmented, corresponding to the described morphological heterogeneity in BC organoids^35,50^. In addition, these hMOs differ in the expression of E-cadherin and GATA-3. E-cadherin is a marker of low metastatic potential and loss of E-cadherin has been associated with increased invasion ^51^, but reduced metastasis and reduced cell proliferation and survival ^52^. In our hands, E-cadherin-negative hMOs displayed high Ki67-determined proliferation rates. Since the effects of E-cadherin loss were attributed to increased reactive oxygen species (ROS) production and TGF-ß signaling ^52^, including the antioxidant N-acetylcysteine and the TGF-ß inhibitors A83-01 and Noggin likely supported proliferation and cell survival of E cadherin low expressing hMOs. Inversely, hMOs with higher expression of E-cadherin demonstrated an intermediate growth rate.

GATA-3 is a transcription regulator required for normal mammary gland development and is associated with stemness and the luminal transcription program ^37^. In recent studies, GATA-3 has been described as a positive prognostic factor linking its expression to a lower Nottingham histologic grade and lower macrophage infiltration ^53^. These results are in line with studies of BC cell lines where GATA-3 reverses epithelial-to-mesenchymal-transition ^54^ and suppresses metastasis ^55^. Here, GATA-3 expressing hMOs were more sensitive towards Birinapant- and Emricasan-induced necroptosis indicating that GATA-3 expression might mark BC that is more sensitive towards PCD, but further research is required to confirm this.

To investigate the susceptibility of the primary hMOs towards PCD induction, we applied Smac mimetics which act as degraders of the IAP E3 ligases, thereby preventing pro-survival ubiquitination on cellular targets and cell death induction ^20^. Both cIAPs and XIAP are reported to be overexpressed in BC ^56^ and are associated with a worse prognosis ^57,58^. Interestingly, whereas cIAP1 was expressed in hMOs derived from all donors, the hMOs from donor #3 displayed basal cIAP2, whereas cIAP2 expression in hMOs from donors #1 and #2 could not be detected. Treatment of the hMOs with Smac mimetics induced loss of viability. hMOs from donor #3 showed a minor reduction of cell viability upon treatment with Birinapant, ASTX-660 and LCL-161, which might be linked to cIAP2 expression compensating for the loss of cIAP1 ^59^. Although suggested as novel treatment options for BC ^56^, in-depth investigations using primary patient-derived model systems were lacking. The strongest influence on cell viability were observed for BV6 and Birinapant, whereas ASTX-660 and LCL-161 showed less potent effects. BV6 and Birinapant are both bivalent IAP inhibitors, whereas the other two Smac mimetics have a monovalent structure ^19^. In the hMOs, bivalent conformations with larger molecular structures have higher efficacies in reducing cell viability ^60^. Importantly, the average pore size of Matrigel has been quantified using scanning electron microscopy (SEM) to 4 µm^2,61^, thereby excluding the matrix as a barrier that prevents the diffusion of bivalent Smac mimetics. Smac mimetics induced caspase-3 cleavage and apoptosis in all tested hMOs. All tested hMOs expressed high levels of endogenous TNF-α which further increased upon incubation with Smac mimetics. This is consistent with the role of IAPs in stabilizing NIK, a potent activator of the non-canonical NF-ĸB pathway and that activates the expression of genes involved in proliferation and cell death, including TNF-α and IAPs themselves ^62^. By doing so, the increased expression levels of TNF-α further drive feed-forward loops that support an inflammatory cellular microenvironment and the induction of apoptosis ^26^.

Lytic forms of PCD, including necroptosis, are promising in overcoming apoptosis resistance observed in many tumors ^63^. Necroptosis is morphologically characterized by cell swelling, membrane rupture and release of highly inflammatory cytokines, chemokines and DAMPs and mediated by the coordinated phosphorylation of receptor-interacting serine/threonine-protein kinase 1 (RIPK1) at S166, RIPK3 at S227 and MLKL at residues T357 and S358 ^40,64^. Activated MLKL induced MLKL oligomerization, plasma membrane accumulation and permeabilization and necroptotic cell death ^64^. Our work demonstrates that apoptosis resistant, primary hMOs are susceptible to Smac mimetic-induced necroptosis and that RIPK1 phosphorylation can be applied as a biomarker for necroptosis in primary BC organoids. Interestingly, many BC cell lines do not express RIPK1, RIPK3 or MLKL and are non-sensitive towards necroptosis. Our findings also indicate expression and phosphorylation of RIPK1, RIPK3 and MLKL upon necroptosis progression in the hMOs, establishing primary, patient-derived organoids as a valuable model for necroptosis in breast cancer. Our findings also reveal that necroptosis in hMOs induces the release of inflammatory mediators, such as IP-10 that, besides danger-associated molecular patterns (DAMPs), like adenosine triphosphate (ATP), lactate dehydrogenase (LDH) and high mobility group box 1 (HMGB1) affect neighboring cells, the cellular microenvironment and infiltration, differentiation and function of immune cells ^65^.

Although LUBAC is well characterized during early stages of TNF-α and Smac mimetic-induced cell death, the roles of linear ubiquitination in controlling necroptosis at the levels of MLKL remains largely unclear. LUBAC inhibition with HOIPIN-8 in the primary hMOs delayed necroptosis onset and lowered cytokine and chemokine secretion. These results agree with our previous study investigating the influence of LUBAC inhibition on necroptosis in the human HT29 colon carcinoma cell line, as well as primary human pancreatic organoids ^34^. These findings might point towards tissue-, cell type- and species-specific modes of necroptosis regulation that further support the use of primary human tumor organoids as experimental models.

Three-dimensional (3D) cell and tissue cultures have emerged as a highly promising tool in cell biology and drug discovery ^66,67^. Multicellular human tumor organoids are driven by adult stem cells and represent a significant advancement in our ability to mimic the intricate functioning of tumors *in vitro*^67–70^. In contrast to conventional 3D spheroids derived from tumor cell lines, primary tumor organoids offer a superior platform due to their unique composition, cultured from primary cells obtained directly from patients and maintained in a genetically stable and proliferative state for extended periods, often up to one year ^67–70^. Contrary to traditional 2D cultures, 3D cell cultures can be cultured in a chemically defined and GMP-compliant manner ^71,72^ and provide more sophisticated and realistic representations of complex human physiology, tissue homeostasis and cell-cell contacts ^66,67^. 3D cell cultures offer a more accurate representation of human physiology at the cellular and tissue level ^66,67^. These cultures enable researchers to recreate the 3D microenvironment, cell-cell interactions and extracellular matrix composition that closely mimic the intricate conditions found in the human body ^66,67^. Consequently, this enhanced physiological relevance makes 3D cell cultures a valuable tool for investigating fundamental biological processes and understanding disease mechanisms with greater precision.

Finally, from an ethical standpoint, the utilization of 3D cell cultures offers a crucial opportunity to reduce the dependence on animal models in pre-clinical research^71,73^. Animal testing has long been a contentious subject due to concerns over animal welfare and the validity of extrapolating results to human systems. By employing human-derived 3D tumor organoids, the need for extensive animal testing can be bypassed, thus minimizing the ethical challenges associated with animal experimentation ^66,67^. Furthermore, this shift allows for a more cost-effective and efficient research approach, as the use of animals can be limited to only the most essential and specialized studies.

Overall, our findings reveal primary hMOs as suitable models to study PCD sensitivities and inflammatory signaling and provide an attractive experimental platform for personalized and precision translational approaches, immunotherapy and drug screening.

## Acknowledgements

The lab of S.J.L.v.W. received funding by the Deutsche Forschungsgemeinschaft (DFG, German Research Foundation) (WI 5171/1-1, WI 5171/4-1, WI 5171/3-2, FU 436/20-1, FU 436/21-1 and project-ID 259130777 – CRC1177), the Deutsche Krebshilfe (70113680), the Wilhelm Sander-Stiftung (2020.008.1), the Frankfurter Stiftung für krebskranke Kinder and the Dr. Eberhard and Hilde Rüdiger Foundation. K.N.W., F.P. and E.H.K.S. acknowledge funding by the iMOL (interfacing image analysis and molecular life-science) DFG Research group (project # 414985841, GRK 2566). F.P. acknowledges funding by the EU Horizon2020 project BRIGHTER (Grant #828931), the EU Horizon-EIC-2021 project B-BRIGHTER (Grant #101057894), and the German Space Agency at DLR (Grant #50W2019 and #50WB2316). The lab of M.B. received funding by DFG (CRC1160 (Project ID 256073931-Z02), CRC/TRR167 (Project ID 259373024-Z01), CRC1453 (Project ID 431984000-S1), CRC1479 (Project ID: 441891347-S1), TRR 359 (Project ID 491676693-Z01), FOR 5476 UcarE (Project ID 493802833-P7)). M.B. and A.G. also acknowledge funding from the German Federal Ministry of Education and Research (BMBF) within the Medical Informatics Funding Scheme PM4Onco–FKZ 01ZZ2322A (M.B.) and EkoEstMed–FKZ 01ZZ2015 (G.A.). We thank the HUB Organoids for their support and the provided organoid lines. We thank Heinz Schewe (Frankfurt Center of Advanced Light Microscopy, Goethe University Frankfurt am Main) for his technical support on the microscopes. The authors thank the research groups of E.H.K.S. and S.J.L.v.W. as well as their former members for advice and discussions.

## Author contributions

K.N.W., F.P. and S.J.L.v.W. designed the experiments and K.N.W. conducted most of the experiments using the hMOs. J.V.S. supported and conducted further experiments with the hMOs. I.K. and A.W. performed the CBA assays and the downstream analysis.

N.D. supported in preliminary testing and provided help in the experiments. F.P. and S.J.L.v.W. supervised the research. E.H.K.S. supported the research. The manuscript was written by K.N.W., F.P. and S.J.L.v.W. All authors revised and agreed on the submitted manuscript.

## Supplementary Figures

**Supplementary Figure 1 related to Figure 3:**
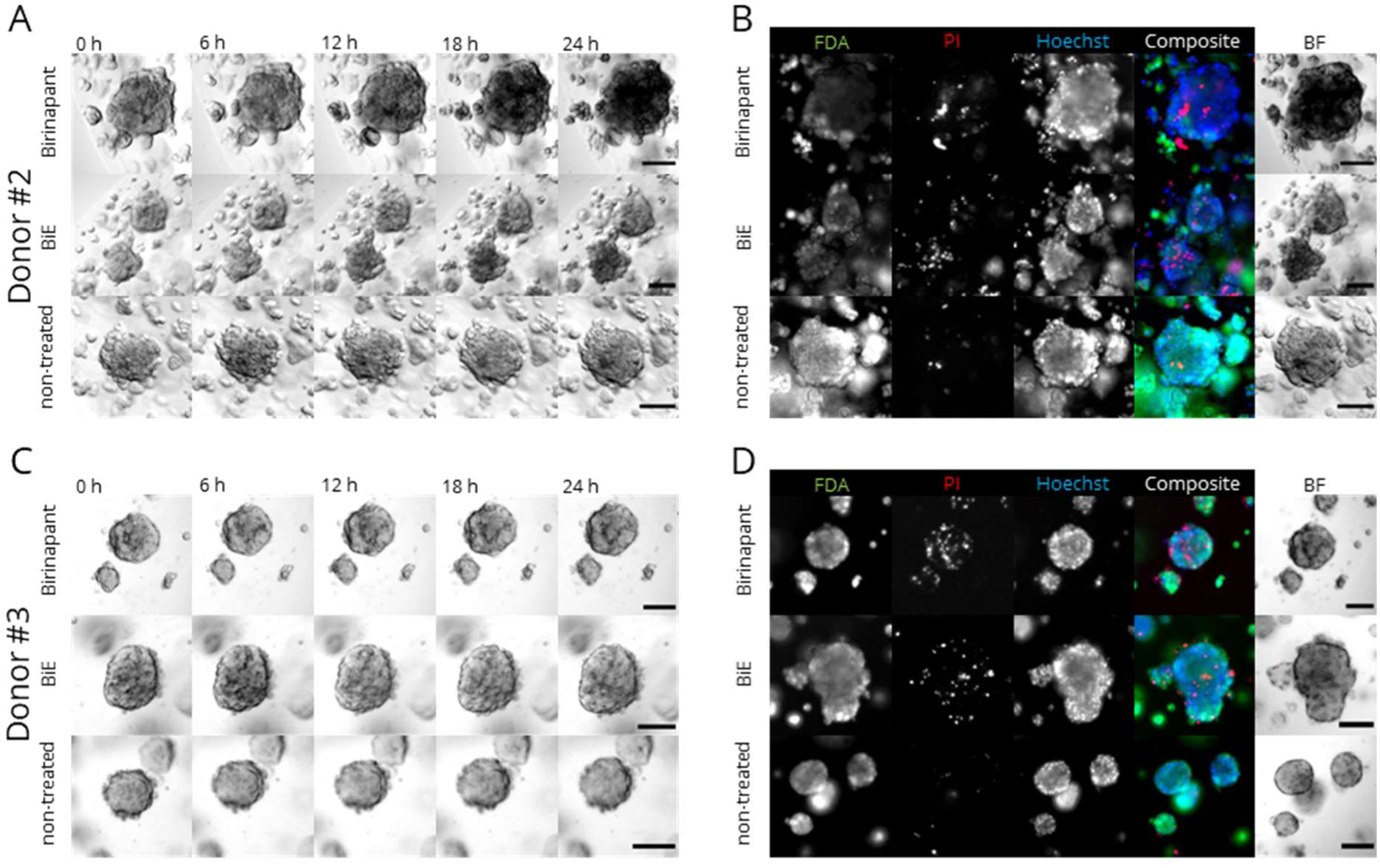
Treatment with Birinapant and in combination with Emricasan lead to cell death in hMOs. (A) hMOs from donor #2 were grown for three to seven days followed by 24 h treatment with 10 µM Birinapant and in combination with 10 µM Emricasan (BiE). Brightfield (BF) time lapse imaging was performed using the Zeiss Z1 Axioimager widefield microscope with 30 min intervals. (B) hMOs from (A) after live imaging were stained using fluorescein-diacetate (FDA, viable cells, green), propidium iodide (PI, dead cells, red) and Hoechst33342 (Hoechst, all nuclei, blue). (C) Same as (A), but hMOs from donor #3 were used. (D) Same as (B), but hMOs from donor #3 were used. Representative images of donor #2 and #3. Scale bars: 100 µm.

**Supplementary Figure 2 related to Figure 3:**
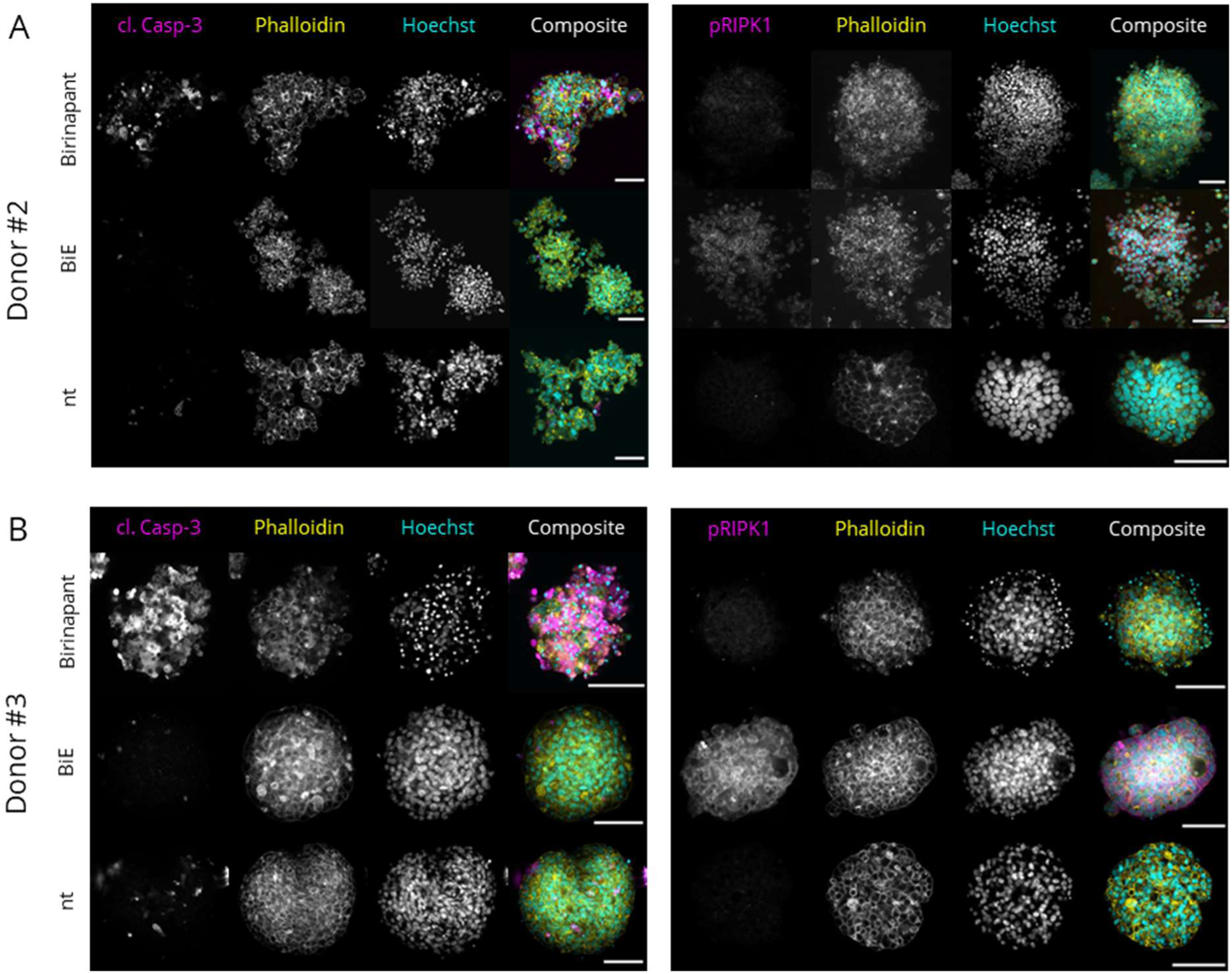
Birinapant- and Emricasan-induced cell death shows hallmarks of apoptosis and necroptosis. (A) hMOs from donor #2 were grown for three to seven days followed by 24 h treatment with 10 µM Birinapant and in combination with 10 µM Emricasan (BiE) were fixed and subjected to immunofluorescence staining against cleaved caspase-3 (cl. Casp-3, magenta) or S166-phosphorylated RIPK1 (pRIPK1, magenta) and counterstained using AF488-phalloidin (yellow) and Hoechst33342 (cyan). Organoids were cleared with CUBIC-2 and imaged using a Zeiss LSM780 confocal microscope. (B) Same as (A), but hMOs from donor #3 were used. Representative images of donor #2 and donor #3. Scale bars: 100 µm.

**Supplementary Figure 3 related to Figure 3:**
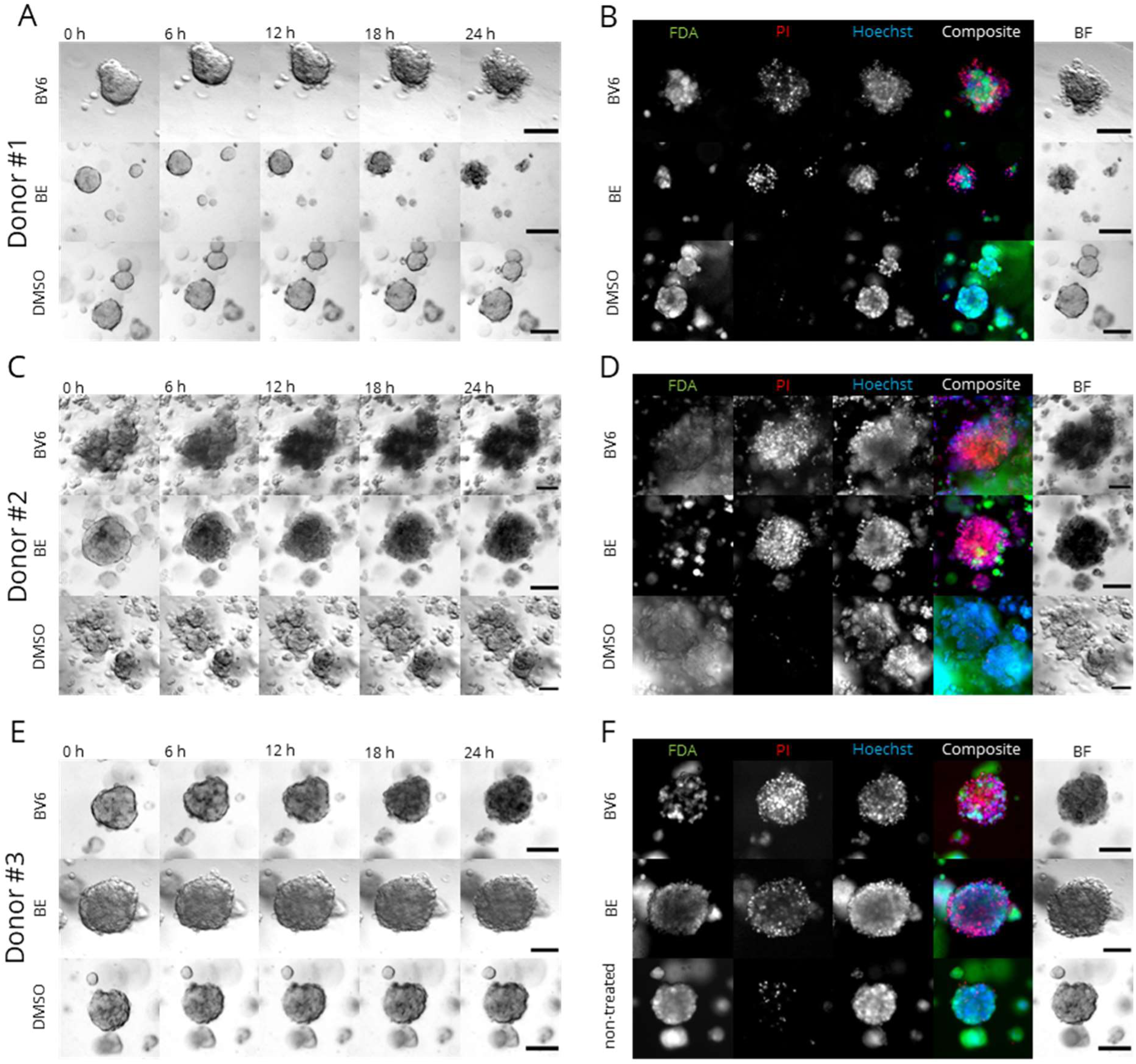
Treatment with BV6 and in combination with Emricasan lead to cell death in hMOs. (A) hMOs from donor #1 were grown for three to seven days followed by 24 h treatment with 10 µM BV6 and in combination with 10 µM Emricasan (BE) or DMSO. Brightfield (BF) time lapse imaging was performed using the Zeiss Z1 Axioimager widefield microscope with 30 min intervals. (B) hMOs from (A) after live imaging were stained using fluorescein-diacetate (FDA, viable cells, green), propidium iodide (PI, dead cells, red) and Hoechst33342 (Hoechst, all nuclei, blue). (C) Same as (A), but hMOs from donor #2 were used. (D) Same as (B), but hMOs from donor #2 were used. (E) Same as (A), but hMOs from donor #3 were used. (F) Same as (B), but hMOs from donor #3 were used. Representative images of hMOs from donors #1, #2 and #3. Scale bars: 100 µm.

**Supplementary Figure 4 related to Figure 3:**
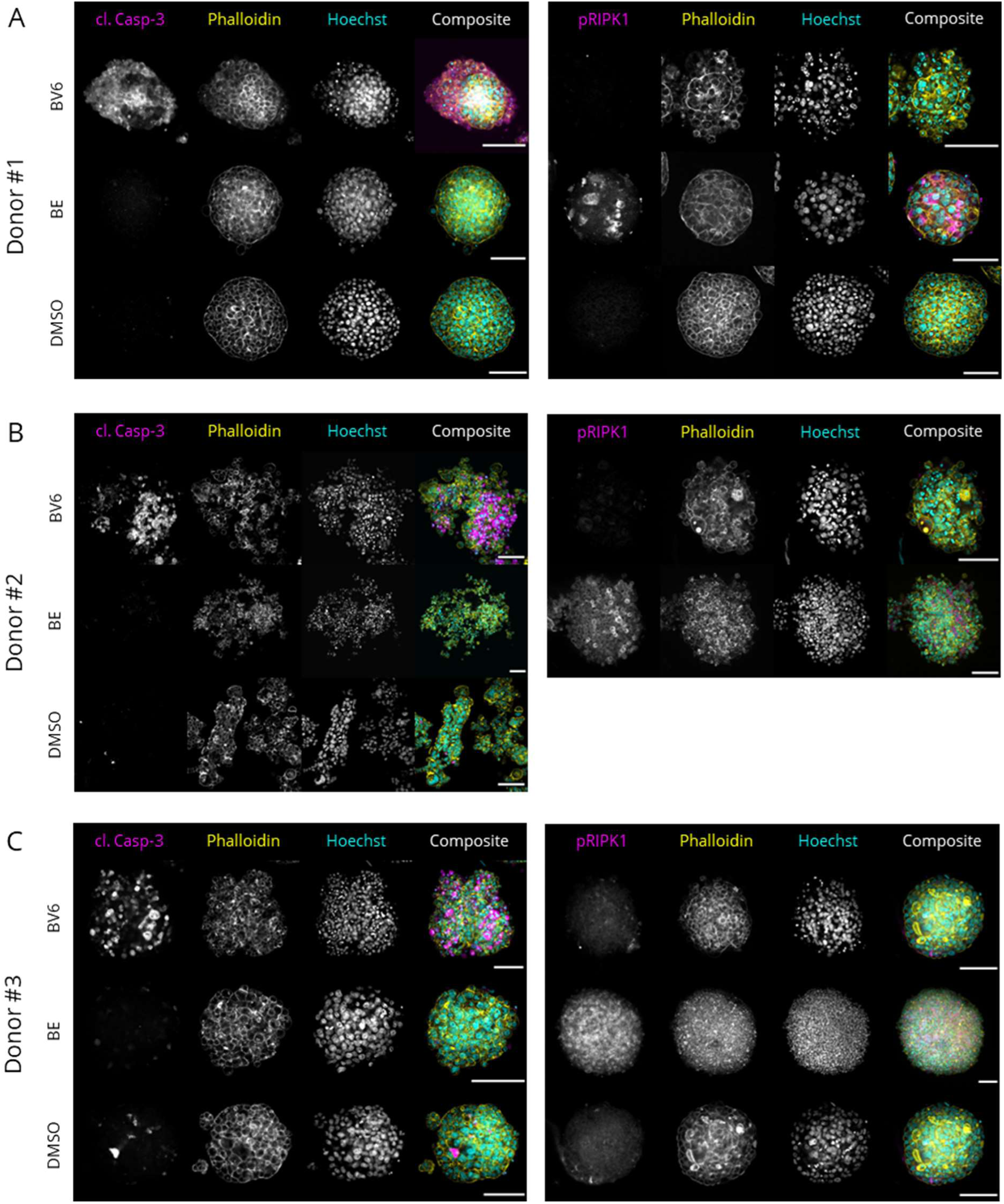
BV6- and Emricasan-induced cell death shows hallmarks of apoptosis and necroptosis. (A) hMOs from donor #1 were grown for three to seven days followed by 24 h treatment with 10 µM BV6 and in combination with 10 µM Emricasan (BE) or DMSO and were fixed and subjected to immunofluorescence staining against cleaved caspase-3 (cl. Casp-3, magenta) or S166-phosphorylated RIPK1 (pRIPK1, magenta) and counterstained using AF488-phalloidin (yellow) and Hoechst33342 (cyan). Organoids were cleared with CUBIC-2 and imaged using a Zeiss LSM780 confocal microscope. (B) Same as (A), but hMOs from donor #2 were used. (C) Same as (A), but hMOs from donor #3 were used. Representative images of hMOs from donors #1, #2 and #3. Scale bars: 100 µm.

**Supplementary Figure 5 related to Figure 4:**
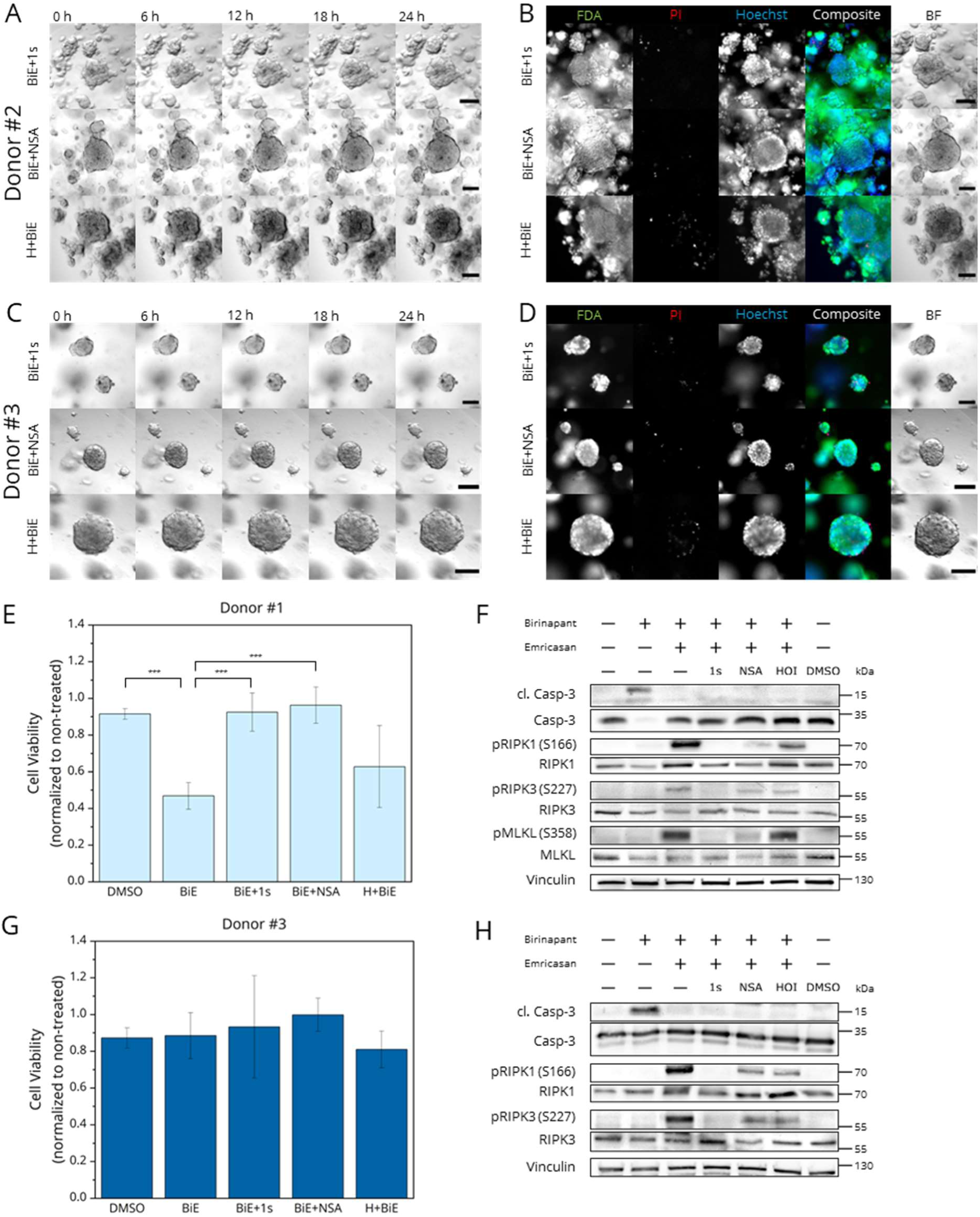
BiE-induced necroptosis is inhibited in hMOs by co-treatment with necroptosis inhibitors and LUBAC inhibitor HOIPIN-8 in different donors. (A) hMOs from donor #2 were grown for three to seven days and pre-treated with 30 µM HOIPIN-8 (H, HOI) for 30 min followed by treatment with 10 µM Birinapant and in combination with 10 µM Emricasan (BiE), 30 µM necrostatin-1s (1s) and 10 µM necrosulfonamide (NSA) as indicated. Brightfield (BF) time lapse imaging was performed using the Zeiss Z1 Axioimager widefield microscope with 30 min intervals. (B) hMOs from (A) after live imaging were stained using fluorescein-diacetate (FDA, viable cells, green), propidium iodide (PI, dead cells, red) and Hoechst33342 (Hoechst, all nuclei, blue). (C) Same as (A), but hMOs from donor #3 were used. (D) Same as (B), but hMOs from donor #3 were used. Scale bars: 100 µm. (E) CellTiter-Glo® viability assays were performed on hMOs from donor #1 treated as in (A). Values were normalized to non-treated controls. Data are shown as mean and standard deviation. *** p < 0.005. (F) hMOs from donor #1, treated as in (A) were harvested and Western blotting was performed with antibodies recognizing cleaved and total caspase-3, phosphorylated and total RIPK1, RIPK3 and MLKL. Vinculin served as loading control. Representative blots of three independent experiments are shown. (G) Same as (E), but hMOs from donor #3 were used. (H) Same as (F), but hMOs from donor #3 were used.

**Supplementary Figure 6 related to Figure 4:**
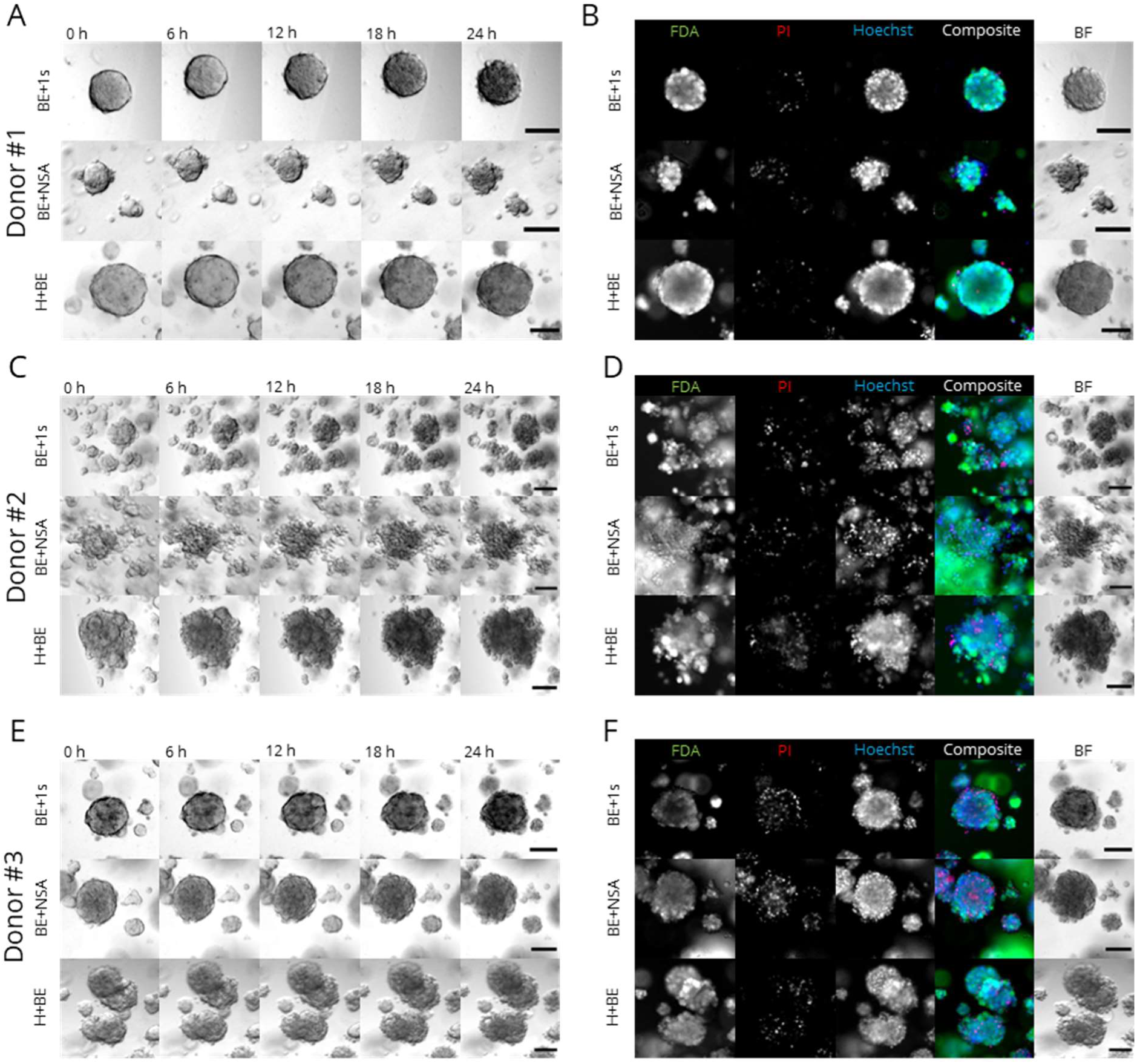
BV6-Emricasan-induced cell death can be inhibited by necroptosis inhibitors and LUBAC inhibitor HOIPIN-8 in hMOs. (A) hMOs from donor #1 were grown for three to seven days and pre-treated with 30 µM HOIPIN-8 (H, HOI) for 30 min followed by treatment with 10 µM Birinapant and in combination with 10 µM Emricasan (BiE), 30 µM necrostatin-1s (1s) and 10 µM necrosulfonamide (NSA) as indicated. Brightfield (BF) time lapse imaging was performed using the Zeiss Z1 Axioimager widefield microscope with 30 min intervals. (B) hMOs from (A) after live imaging were stained using fluorescein-diacetate (FDA, viable cells, green), propidium iodide (PI, dead cells, red) and Hoechst33342 (Hoechst, all nuclei, blue). (C & D) Same as (A & B), but hMOs from donor #2 were used. (E & F) Same as (A & B), but hMOs from donor #3 were used. Scale bars: 100 µm.

**Supplementary Figure 7 related to Figure 4:**
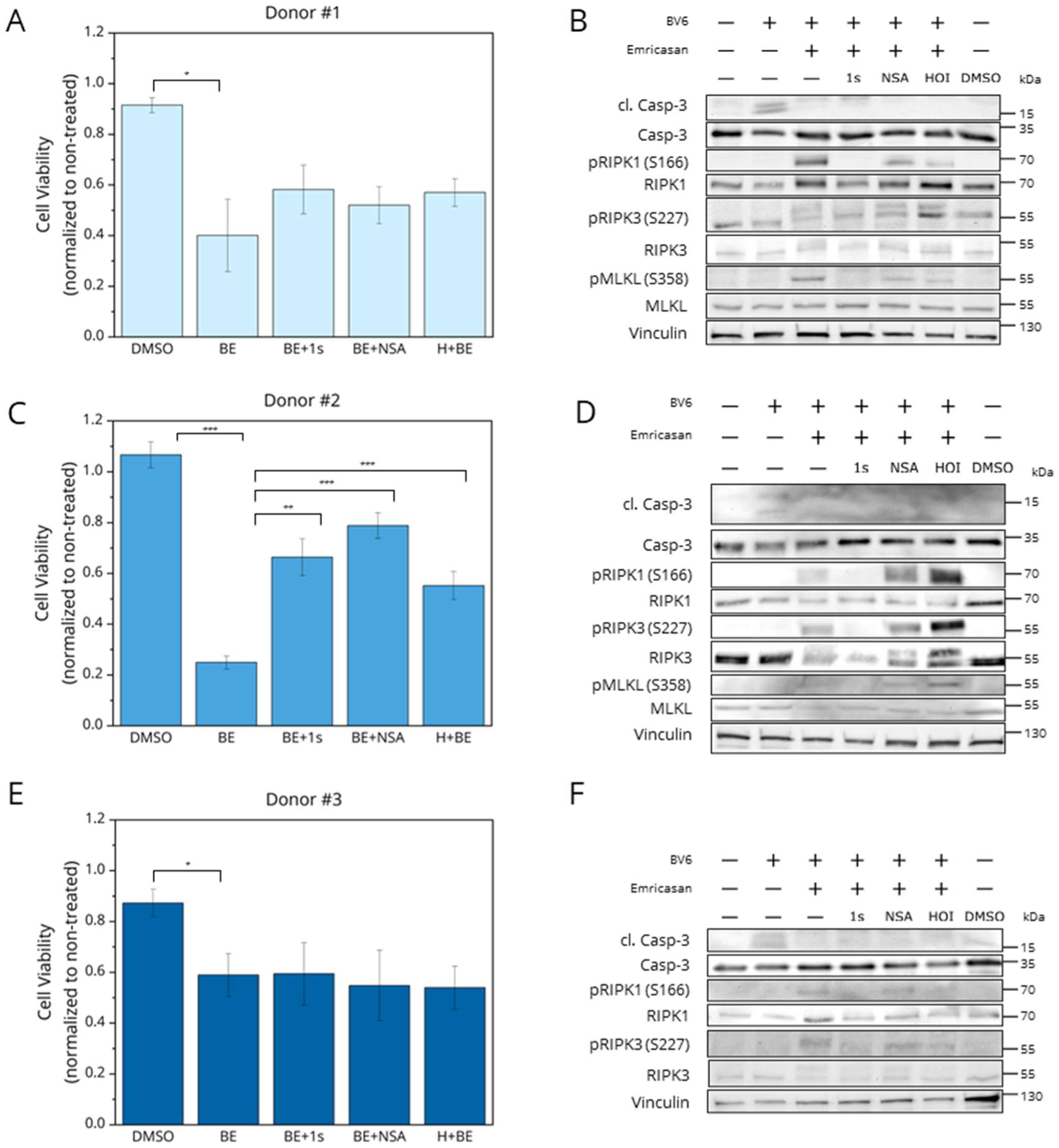
Necroptosis induction can be inhibited by RIPK1 inhibitor Nec-1s, MLKL inhibitor NSA and LUBAC inhibitor HOIPIN-8. (A) hMOs from donor #1 were grown for three to seven days and pre-treated with 30 µM HOIPIN-8 (H, HOI) for 30 min followed by treatment with 10 µM Birinapant and in combination with 10 µM Emricasan (BiE), 30 µM necrostatin-1s (1s) and 10 µM necrosulfonamide (NSA) as indicated followed by CellTiter-Glo® viability assays. Values were normalized to non-treated controls. Data are shown as mean and standard deviation. (B) hMOs from donor #1, treated as in (A) were harvested and Western blotting was performed with antibodies recognizing cleaved and total caspase-3, phosphorylated and total RIPK1, RIPK3 and MLKL. Vinculin served as loading control. Representative blots of three independent experiments are shown. (C & D) Same as (A & B), but hMOs from donor #2 were used. (E & F) Same as (A & B), but hMOs from donor #3 were used. * p < 0.05, ** p< 0.01, *** p < 0.005.

**Supplementary Figure 8 related to Figure 5:**
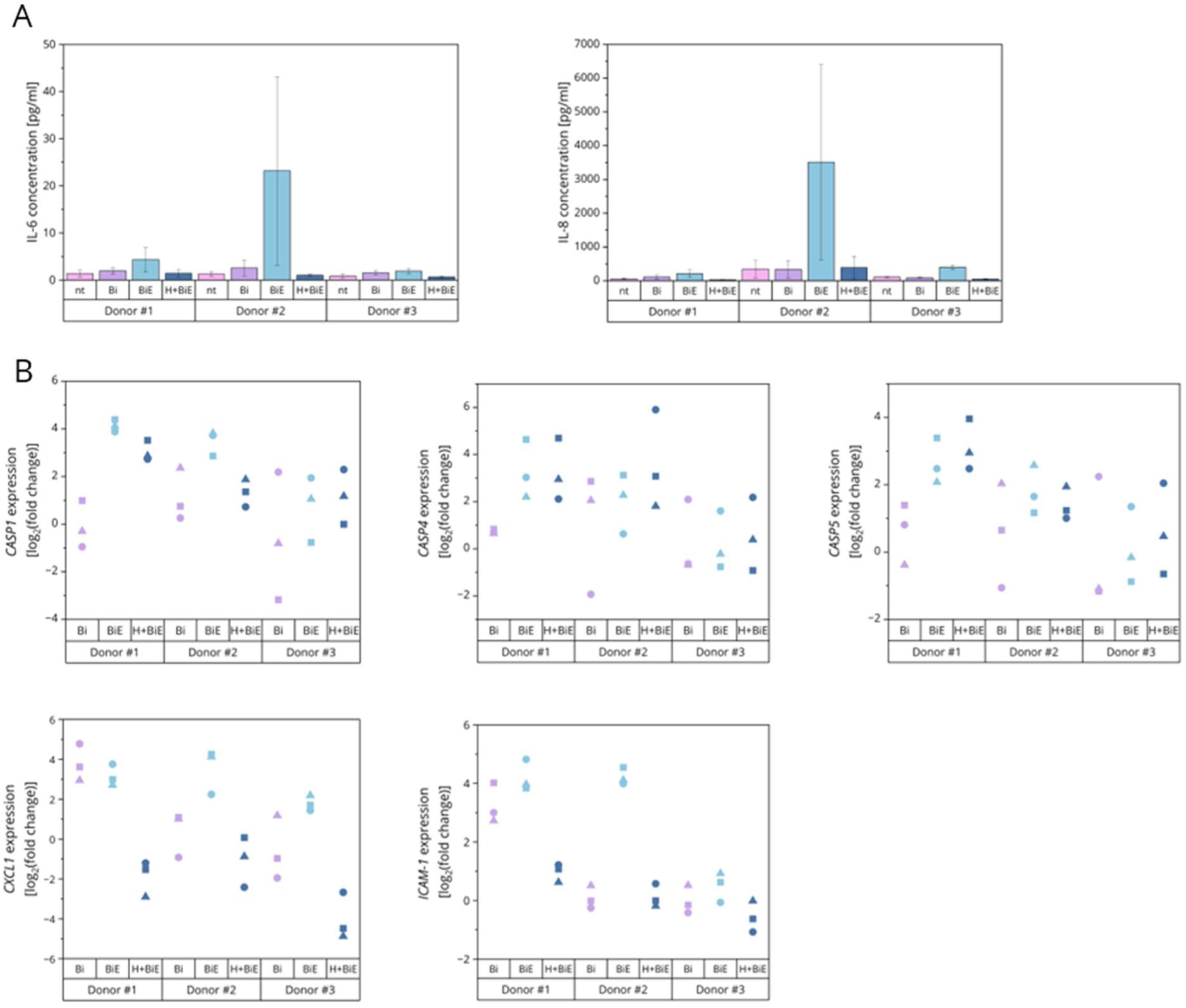
Necroptosis induces inflammatory signaling that can be reversed by co-treatment with HOIPIN-8 in hMOs. (A) FACS-based CBA assays to quantify secretion of IL-6 and IL-8 in the indicated hMOs grown for three to seven days and then pre-treated with 30 µM HOIPIN-8 for 30 min followed by treatment with 10 µM Birinapant and in combination with 10 µM Emricasan (BiE). Data are shown as mean with standard deviation and are from three independent experiments. (B) mRNA expression levels of *CASP1*, *CASP4*, *CASP5*, *CXCL1* and *ICAM1* in the indicated hMOs grown for three to seven days, pre-treated with 30 µM HOIPIN-8 for 30 min followed by treatment with 10 µM Birinapant and imcombination with 10 µM Emricasan (BiE). Gene expression was normalized against non-treated conditions and *RPII*, *18S*, *TBP* and *RPL13* mRNA expression and is presented as log_2_(fold change).

**Supplementary Figure 9 related to Figure 5:**
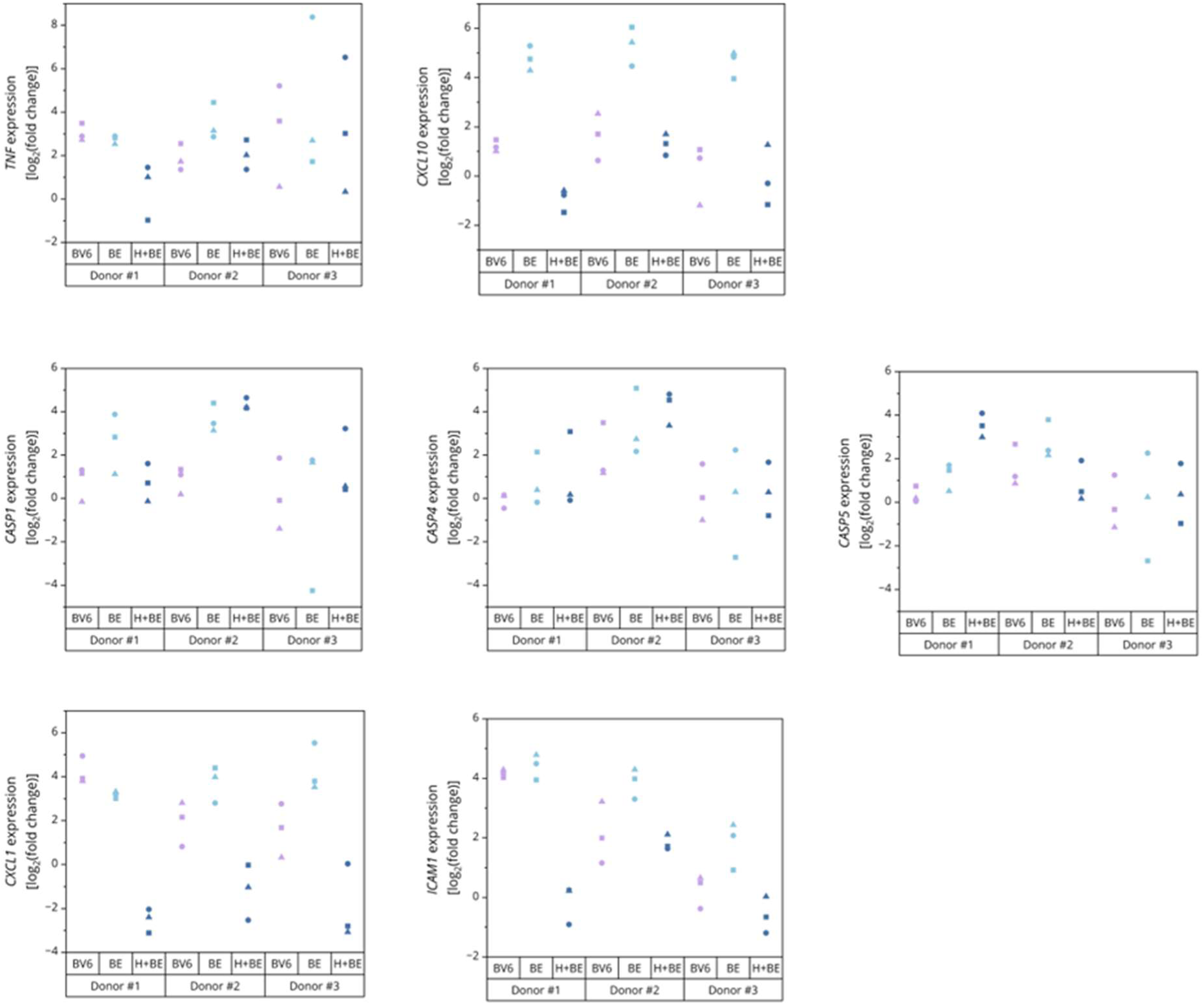
BV6-Emricasan-induced necroptosis causes inflammatory signaling that can be reversed by co-treatment with HOIPIN-8 in hMOs. mRNA expression levels of *TNF, CXCL10*, *CASP1*, *CASP4*, *CASP5*, *CXCL1* and *ICAM1* in the indicated hMOs grown for three to seven days, pre-treated with 30 µM HOIPIN-8 for 30 min followed by treatment with 10 µM BV6 (B) with and without 10 µM Emricasan (E). Gene expression was normalized against non-treated conditions and *RPII*, *18S*, *TBP* and *RPL13* mRNA expression and is presented as log_2_(fold change).

